# SpatialCTD: a large-scale TME spatial transcriptomic dataset to evaluate cell type deconvolution for immuno-oncology

**DOI:** 10.1101/2023.04.11.536333

**Authors:** Jiayuan Ding, Julian Venegas, Qiaolin Lu, Yixin Wang, Lidan Wu, Wei Jin, Hongzhi Wen, Renming Liu, Wenzhuo Tang, Zhaoheng Li, Wangyang Zuo, Yi Chang, Yu Leo Lei, Patrick Danaher, Yuying Xie, Jiliang Tang

**Affiliations:** Department of Computer Science and Engineering, Michigan State University, East Lansing, USA; Department of Computational Mathematics, Science and Engineering, Michigan State University, East Lansing, USA; Department of Statistics and Probability, Michigan State University, East Lansing, USA; School of Artificial Intelligence, Jilin University, Changchun, China; Department of Bioengineering, Stanford University, Palo Alto, USA; Department of Biostatistics, University of Washington, Seattle, USA; Department of Computer Science, Zhejiang University of Technology, Hangzhou, China; NanoString Technologies, Seattle, USA; Department of Periodontics and Oral Medicine, University of Michigan School of Dentistry, Ann Arbor, USA; University of Michigan Rogel Cancer Center, Ann Arbor, USA

**Keywords:** Cell type deconvolution, Spatial transcriptomic data, Benchmark dataset, Tumor microenvironment, Human immuno-oncology, Graph Neural Networks

## Abstract

Recent technological advancements have enabled spatially resolved transcriptomic profiling but at multi-cellular resolution. The task of cell type deconvolution has been introduced to disentangle discrete cell types from such multi-cellular spots. However, existing datasets for cell type deconvolution are limited in scale, predominantly encompassing data on mice, and are not designed for human immuno-oncology. In order to overcome these limitations and promote comprehensive investigation of cell type deconvolution for human immuno-oncology, we introduce a large-scale spatial transcriptomic dataset named SpatialCTD, encompassing 1.8 million cells from the human tumor microenvironment across the lung, kidney, and liver. Distinct from existing approaches that primarily depend on single-cell RNA sequencing data as a reference without incorporating spatial information, we introduce Graph Neural Network-based method (i.e., GNNDeconvolver) that effectively utilize the spatial information from reference samples, and extensive experiments show that GNNDeconvolver often outperforms existing state-of-the-art methods by a substantial margin, without requiring single-cell RNA-seq data. To enable comprehensive evaluations on spatial transcriptomics data from flexible protocols, we provide an online tool capable of converting spatial transcriptomic data from other platforms (e.g., 10x Visium, MERFISH and sci-Space) into pseudo spots, featuring adjustable spot size. The SpatialCTD dataset and GNNDeconvolver implementation are available at https://github.com/OmicsML/SpatialCTD, and the online converter tool can be accessed at https://omicsml.github.io/SpatialCTD/.

## 1 Introduction

Spatial transcriptomics has provided novel opportunities to gain a deeper understanding of spatial heterogeneity of tissue architecture [9, 14], analyze cellular interactions [15, 23], and investigate tissue neighborhoods and local features that contribute to disease [14, 24]. However, the majority of existing relatively affordable spatial transcriptomic technologies, such as Spatial Trascriptomics [19], 10X Visium [1], and Slide-seq [17, 20], detect expressed RNAs at the spot-level in space where each spot contains multiple cells, resulting in a mixed gene expression profile that does not provide spatial singlecell resolution. This limitation hinders the accurate quantification of spatial cellular distribution and subsequent analysis.

To overcome the aforementioned limitation, the task of cell type deconvolution has been introduced to decompose spatial mixture gene expression into cellular proportions per spot. Recent years have witnessed an increasing number of computational approaches for cell type deconvolution. For example, Seurat [21] and Giotto [7] leverage the enrichment score of a gene set (such as cell-type-specific marker genes obtained from scRNA-seq data) to estimate the likelihood of each cell type in a given spot. SPOTlight [8] utilizes seeded non-negative matrix factorization (NMF) regression to deconvolve spatial transcriptomics spots with single-cell transcriptomes. Stereoscope [5] develops a Bayesian model that integrates information from both single-cell and spatial transcriptomics data to estimate the probability of each cell type at each location within the tissue sample. Cell2location [12] is a Bayesian hierarchical model to analyze spatial expression counts with a spatially informed prior on cell-type compositions for cell type composition inference. In addition, deep learning based methods have also been developed to advance cell type devconvolution. Tangram [6] employs a deep learning framework to discover a spatial alignment for scRNA-seq data, and then utilizes the identified spatially co-expressed gene modules from the alignment process to deduce the existence of various cell types within a given tissue sample. DSTG [18] utilizes graph-based convolutional networks to estimate cell type proportion of real spatial transcriptomics data from pseudo spatial transcriptomics with established cell type compositions for reference.

However, the majority of existing datasets to study cell type deconvolution are relatively small, unrelated to human tissues, and not specifically designed for immuno-oncology purpose, which is to combat neoplasms through the investigation of interactions between tumor cells and the immune system. For example, Lulu Yan et al. [25] propose three benchmark datasets about mouse tissues for benchmarking deconvoluting spatial transcriptomic data, and the sum of the three datasets are as small as 80,000 cells. Bin Li et al. [13] simulate 32 datasets for cell type deconvolution from the scRNA-seq data as the ground truth, which may not fully capture the biological variability in real tissues and reflect the complexity and heterogeneity of real biological systems. Note that the composition and density of immune cells in the tumor microenvironment (TME) profoundly influence tumor progression and success of anti-cancer therapies [22]. Thus, to facilitate the cell type deconvolution research, a spatial transcriptomic dataset in the tumor microenvironment for human immuno-oncology purpose is a pressing need.

To bridge this gap, we introduce a large-scale spatial transcriptomic dataset from human tumor microenvironment with *1*.*8 millions cells* to evaluate cell type deconvolution for immuno-oncology. Different from existing methods that predominantly rely on single-cell RNA sequencing data as a reference without spatial information, we introduce Graph Neural Network-based method i.e., GNNDeconvolver, which effectively leverage the spatial information of reference samples with known cell compositions to infer unlabeled ones. Extensive experiments demonstrate that GNNDeconvolver usually outperforms existing state-of-the-art methods by a large margin. Note that GNNDeconvolver is a reference-free approach, which circumvents the need for utilizing single-cell RNA-seq data as a reference. To enable comprehensive evaluations for cell type deconvolution with spatial transcriptomics data from diverse protocols, we provide an online tool to convert spatial transciptomic data at a single-cell resolution from other platforms (e.g., 10x Visium, MERFISH and sci-Space) to pseudo spots with adjustable spot size.

## 2 Results

### 2.1 An overview of our generated SpatialCTD dataset

The majority of current benchmark datasets for cell type deconvolution are primarily composed of mouse data [4, 13, 25], simulated from single-cell RNA sequencing data instead of real transcriptomic data [13, 25], and not derived from the human tumor microenvironment [2–4, 8, 13, 25]. To address the dearth of benchmark datasets for cell type deconvolution in human immuno-oncology research, we introduce SpatialCTD, a large-scale human-based spatial transcriptomic dataset. The design purpose of SpatialCTD is to enable flexible and comprehensive cell type deconvolution research in immunooncology. We initially collect spatial transcriptomic data at a single-cell resolution from CosMx platform [10] using the spatial molecular imaging technique. The collected datasets encompass approximately 2 million cells, derive from 20 distinct samples originating from human lung, kidney, and liver tissues, and are captured at a single-cell resolution. Given that the collected data is at a single-cell resolution with known single-cell cell type, a grid-based approach is employed to generate pseudo spots from the spatial transcriptomic data, allowing for cell type deconvolution at spot level while maintaining the integrity of the true cellular microenvironment and biological variance(Fig. 1). Regarding the cell type of each individual cell, we directly use the cell type annotation from the original article. To simulate the transcriptome profile of each pseudo spot, the expression profiles of all cells located within one grid region are summed, with the coordinates of the simulated spot set to the center of the region. The ground truth for each spot is determined by calculating the percentage of cell types within the corresponding spot. The generated benchmarking dataset called SpatialCTD presently comprises three human tissues including human lung, kidney and liver. From each sample, SpatialCTD encompasses three files: a spot gene expression file indicating mixed gene expression of multiple cells at the spot level, a spot location file indicating the spatial location of spots within the tissue, and a ground truth file to reveal the cell type proportion of generated pseudo spots (Fig. 1). More details about how to generate SpatialCTD can be referred to Section 4.

**Figure 1:**
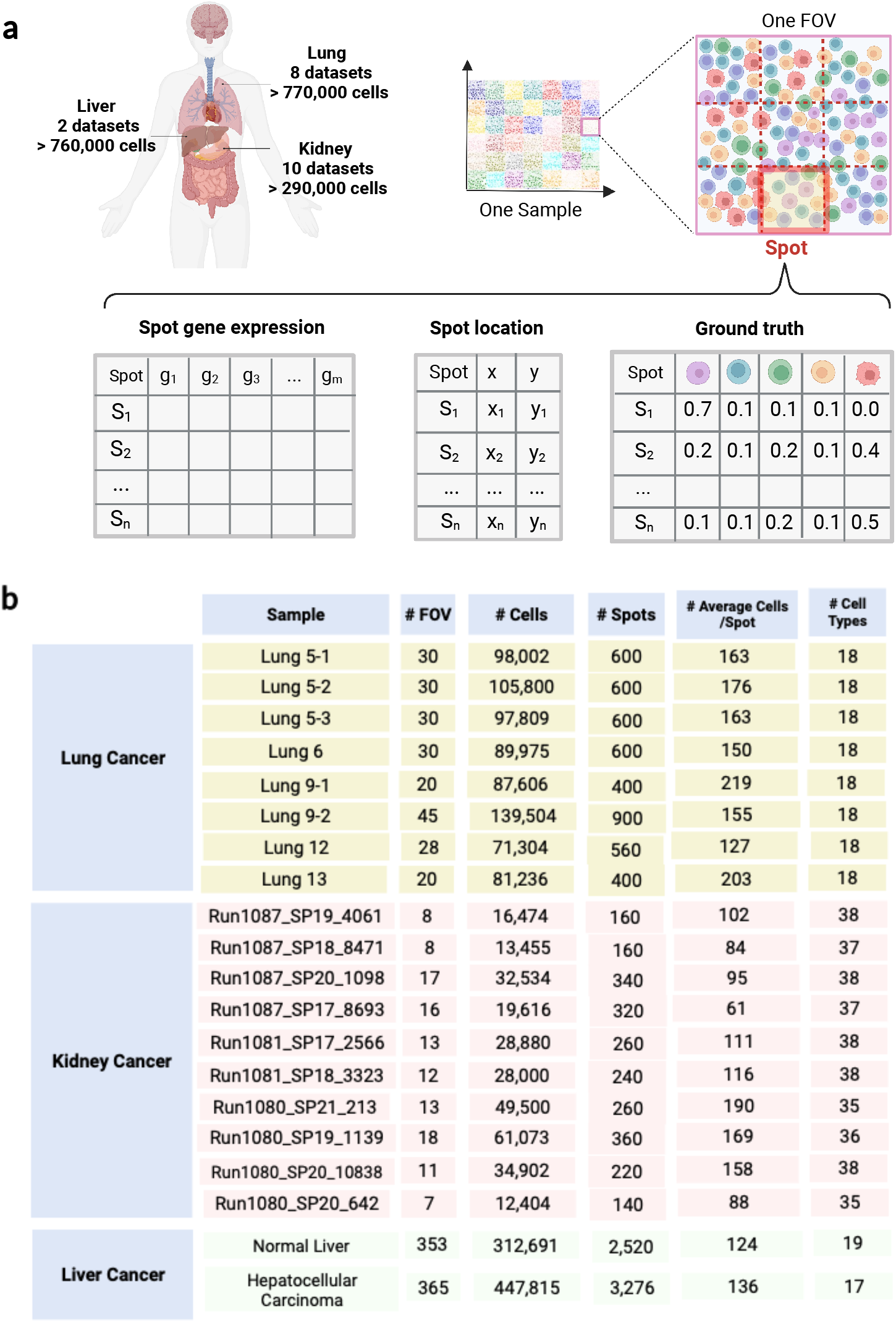
An Overview of SpatialCTD. (**a**), The method for generating the SpatialCTD dataset. SpatialCTD comprises three distinct human tissues, namely, lung, kidney, and liver. For each sample in tissues, SpatialCTD consists of a spot gene expression file, a spot location file, and a ground truth file. (**b**), A summary of SpatialCTD.

In SpatialCTD, a total of 20 samples are included. Of these, 8 are obtained from human lung tissue (Fig. 1). Specifically, **Lung 5-1, 5-2**, and **5-3** are from the same patient, while **Lung 9-1** and **9-2** are from another patient. The remaining lung tissue samples are collected from different patients. Regarding the human kidney tissue, samples of **Run1087_SP19_4061,_Run1087_SP20_1098**, and **Run1080_SP20_10838** are from the one patient, while the remaining kidney tissue samples are obtained from different patients. For the human liver tissue, the normal sample is collected from a 35-year-old male patient, while the hepatocellular carcinoma sample is from a 65-year-old female patient. The field of view (FOV) layout of samples in lung (Fig. 7 in Supplementary) and liver tissues (Fig. 9 in Supplementary) is highly regular, whereas the FOV layout of samples in kidney tissue (Fig. 8 in Supplementary) is less regular. Each tissue has more than 17 cell types, resulting in a highly detailed cell type resolution. The spot size is determined based on the mean and median spot size from the GeoMx platform, which are 37456.28 *μm*^2^ and 24168.74 *μm*^2^, respectively. For lung and kidney tissues, each field of view (FOV) is divided into 20 (4*5) pseudo spots, leading to a spot area of 32338.2067 *μm*^2^. Similarly, for human liver tissue, each FOV is divided into 9 (3*3) pseudo spots, bringing about a spot area of 28709.9136 *μm*^2^. Both of these spot sizes fall within a reasonable range. The average number of cells per spot ranges from 61 to 219. After filtering out spots with low quality, a total of 771,236 cells, 296,838 cells, and 760,506 cells are retained for human lung, kidney, and liver tissues, respectively. To the best of our knowledge, this is the first large-scale dataset of real spatial transcriptomics across several human tissues from tumor microenvironment, which enable advanced and extensive cell type deconvolution research for immuno-oncology.

### 2.2 Comparison between existing benchmark datasets and SpatialCTD

To demonstrate the necessity of the SpatialCTD dataset, we compare, from both biological variance and statistical perspectives, SpatialCTD with existing simulated cell type deconvolution benchmark datasets.

#### Biological variance

We first illustrate the presence of cellular heterogeneity in SpatialCTD collected from the tumor microenvironment (TME), as well as in the simulated spatial transcriptomic data derived from single-cell RNA sequencing data in Experiment 1, and then reveal the existence of cellular heterogeneity in SpatialCTD between normal and tumor tissues in Experiment 2.

In Experiment 1, we initially extract a particular cell type from our dataset (SpatialCTD Lung 5-1) and one simulated dataset (D7 [13]). D7 is obtained from single-cell RNA sequencing data. Note that both SpatialCTD and D7 pertain to the human lung tissue. As a demonstration, the macrophage cell population is collected, comprising 7,405 cells from Lung 5-1 and 2,406 cells from D7. We obtain 942 common genes from two datasets and then perform differential expression (DE) analysis using the R package limma [16], setting the adjusted P value to less than 0.05, between two groups of cells. Subsequently, we identify 283 and 101 DE genes from the DE analysis in macrophage and fibroblast cells, respectively. The same phenomenon can be observed in other major cell types, like B cells, CD4 cells, CD8 cells, endothelial cells and mast cells (Fig. 14 in Supplementary). Experiment 1 provides evidence that cellular heterogeneity is present even for the identical cell type in SpatialCTD compared to D7. SpatialCTD represents the authentic spatial transcriptomic data and is more representative of the actual biological system.

In Experiment 2, we conduct the same DE analysis above between SpatialCTD samples of healthy liver tissue and hepatocellular carcinoma to explore if cellular heterogeneity exists between normal and malignant tissues in spatial transcriptomic data. We use inflammatory macrophage as an example. Healthy liver tissue and hepatocellular cancer yield 5,882 and 14,797 cells, respectively, out of 332,877 and 460,441 cells. The DE genes are then found via DE analysis. TTR, APOA1, HSPA1A, S100A9, S100A8, and SERPINA1 are strongly expressed in the healthy liver tissue but not in hepatocellular carcinoma, whereas HLA-A, CD74, GPX1, HLA-DRB1, PSAP, and MALAT1 are substantially expressed in hepatocellular carcinoma but not in the healthy liver tissue (Fig. 15 and Fig. 16 in Supplementary). For erythroid cells, TTR, APOA1, HSPA1A, APOC1 and SERPINA1 are highly expressed in the healthy liver tissue but lowly expressed in hepatocellular carcinoma while HLA-A, MALAT1, CD74 and B2M are highly expressed in hepatocellular carcinoma but lowly expressed in the healthy liver tissue. The same phenomenon can be observed in other major cell types, like B cells, endothelial cells, T cells and stellate cells (Fig. 15 and Fig. 16 in Supplementary). The two preceding investigations demonstrate the existence of cellular heterogeneity in both real and simulated spatial transcriptomic data, as well as in normal and cancerous tissues. This underscores the need of SpatialCTD to evaluate immuno-oncology cell type deconvolution.

#### Statistical difference

We conduct a thorough comparison of SpatialCTD with a number of existing cell type deconvolution benchmark datasets (Fig. 2). Mouse Embryo [25], MPOA [25], Mouse Cortex [13], and Mouse Visual Cortex [13], adopted a similar approach as ours to generate pseudo spots from spatial transcriptomics (ST) data at the single-cell level. However, these studies were conducted solely on mice and did not involve human samples. Additionally, Mouse Brain [25], Simulated ST for Human Lung [13], and Simulated ST for Human Liver [13] were simulated from single-cell RNA sequencing data rather than actual ST data, and were not derived from TME tissues. Similarly, Mouse Posterior Brain [4] and Mouse Olfactory Bulb [4] are mice based. Despite HEK293T, CCRF-CEM [2] and Human PDAC [3] derived from human tissue, they do not originate from the tumor microenvironment. Furthermore, it is worth noting that the datasets under comparison have a limited sample size, with the largest comprising only 60,000 cells, whereas SpatialCTD comprises up to 770,000 cells within a single tissue. In addition, SpatialCTD presents several distinctive features that are not yet commonly available in existing benchmark datasets, including spot images, which offer a complementary modality for the cell type deconvolution task. Additionally, SpatialCTD provides spatial location data for both cellular and subcellular entities, which has the potential to prompt further cutting-edge research utilizing these features. For example, the identification of cell states in a tissue can be facilitated by cellular location within the tissue. Changes in the spatial distribution of cells within a tissue have been linked to alterations in cell type composition and distribution, which are often associated with disease. The cell and subcellular location can be also further explored to aid in the investigation of ligand-receptor interactions. SpatialCTD is the first large-scale dataset of real spatial transcriptomics collected from various human tissues within the tumor microenvironment. This unique dataset provides an unprecedented opportunity for studying cell type deconvolution in the field of immuno-oncology.

**Figure 2:**
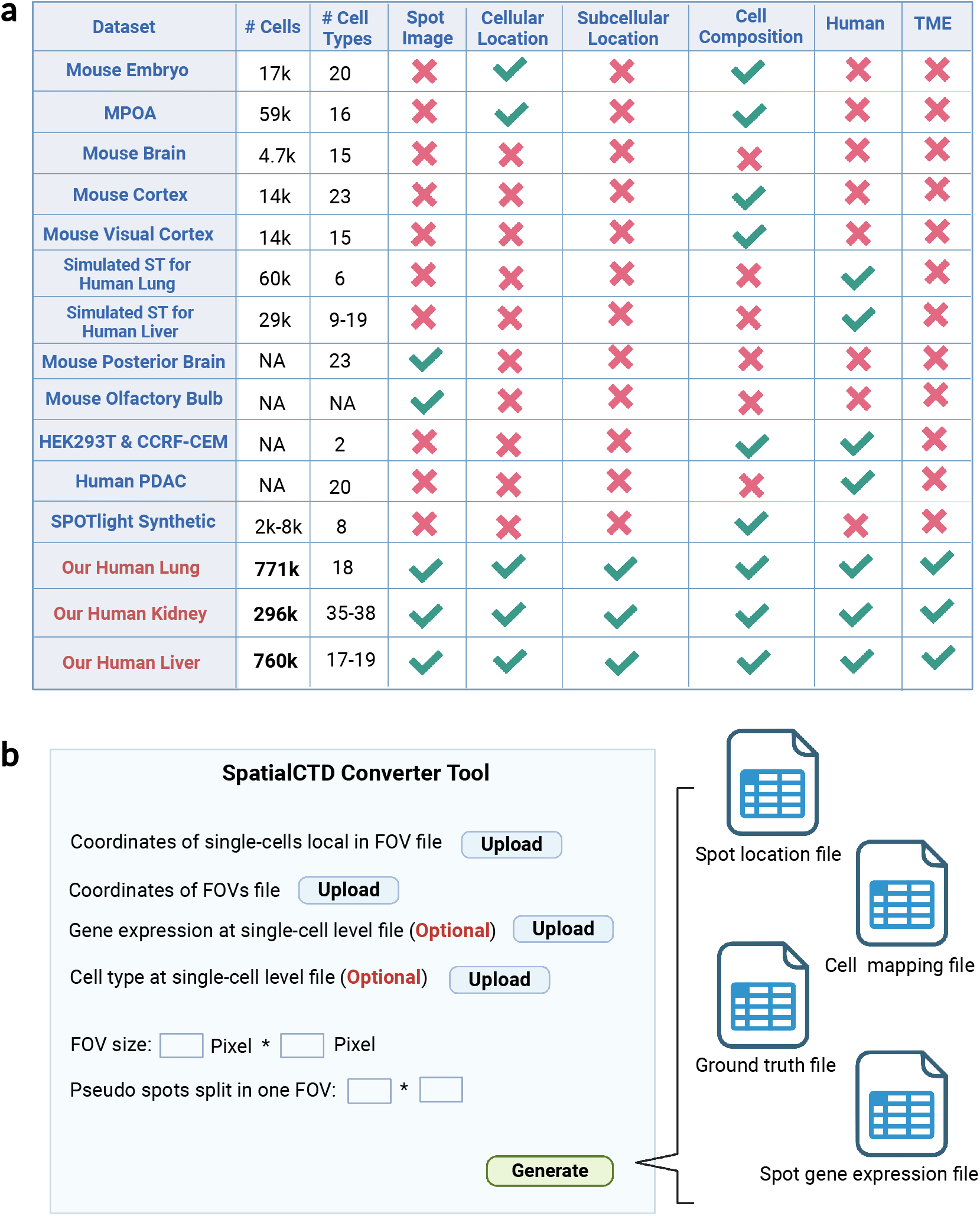
Comparison between SpatialCTD and existing cell type deconvolution benchmark datasets, and SpatialCTD converter tool. **a**, Statistical comparison between SpatialCTD and existing cell type deconvolution benchmark datasets. **b**, An interface of the web-based SpatialCTD converter tool to convert spatial transcriptomic data at a single-cell resolution to ST pseudo spots for cell type deconvolution benchmarking.

### 2.3 Online conversion tool

To ease the process of converting spatial transcriptomics data at single-cell resolution to pseudo spots for benchmarking cell type deconvolution, we have developed a web-based online conversion tool. We provide a friendly web interface for researchers to upload four files, with the first two being mandatory and the latter two being optional. The first file (coordinates of single-cell local in FOV file) contains single-cell level coordinates local in field of view (FOV) (Fig. 2). The second file (coordinates of FOV file) indicates the coordinates of FOV in the slide, which is the spatial coordinates of the center point of each FOV. The third file (single-cell level gene expression file) has gene expression at the single-cell level, which is optional, and the fourth file (single-cell level cell type file) includes single-cell level cell type label, which is also optional. Researchers can refer to the sample data file on our GitHub repository for the specific column naming of each file. Furthermore, researchers need to specify the pixel size of each FOV. We assume that the size of each FOV in the slide is identical. Note that we allow researchers to customize the size of the pseudo spot by specifying how the spots in the FOV are divided.

Upon uploading the aforementioned files and providing the necessary information, researchers can generate the pseudo spot related files by clicking the ‘Generate’ button. The output files include the spot location file, which contains the spatial information of each pseudo spot, and the cell mapping file, which displays the mapping relationship between the pseudo spot ID and the cell ID. The ground truth file has the cell type proportion per spot, and the spot gene expression file provides spot-level gene expression data. If the researcher only uploads the first two files, the tool will only generate the spot location file and the cell mapping file. Researchers can generate the ground truth file and the spot gene expression file offline using the cell mapping file. We anticipate that SpatialCTD will expand beyond the three human tissues we have currently generated, and thus, we hope that developing such tool can simplify the conversion process of other spatial transcriptomics data.

### 2.4 Performance Comparison in SpatialCTD

In this subsection, we design GNNDeconvolver as a Graph Neural Network-based approach that serves as a robust and strong baseline method for analyzing SpatialCTD. In this study, we assume that cell composition labels are either present for the entire sample or not present at all. Prior to applying GNNDeconvolver, we construct a linked graph that connects labeled and unlabeled nodes where labeled nodes denote spots with established cellular compositions in reference samples, and unlabeled nodes correspond to spots within the query sample to be inferred for the cell type proportion. Specifically, we treat each spot as a node in the graph, and build up an internal graph for each sample based on spatial distance (euclidean distance) and the similarity of gene expression between node pairs. We then establish connections between pairs of nodes across different graphs based on the similarity of gene expression. This approach enables the connection of both labeled and unlabeled nodes in the graph. For example, we have three samples: sample 1, sample 2, and sample 3. Sample 1 and sample 2 are labeled with cell type proportion for each spot or node, and hence are referred to as reference or training data. Sample 3, on the other hand, is the test or query data that requires prediction by the model. To achieve this goal, we construct a graph by linking nodes in sample 3 with those in sample 1 and sample 2 that exhibit similar gene expression (Fig. 3). Our approach is based on the assumption that nodes with similar gene expression are likely to have similar cell type proportions. We then apply GNNDeconvolver to the constructed graph. GNNDeconvolver incorporates two graph convolutional layers [11], followed by a softmax function that produces the cell type composition from the output logits. The fundamental principle of GNNDeconvolver is to utilize the graph convolutional operator to propagate the information cross samples and nodes, and thus enable the prediction of cell composition for the unlabeled node. More details about GNNDeconvolver can be referred to Section 4.

**Figure 3:**
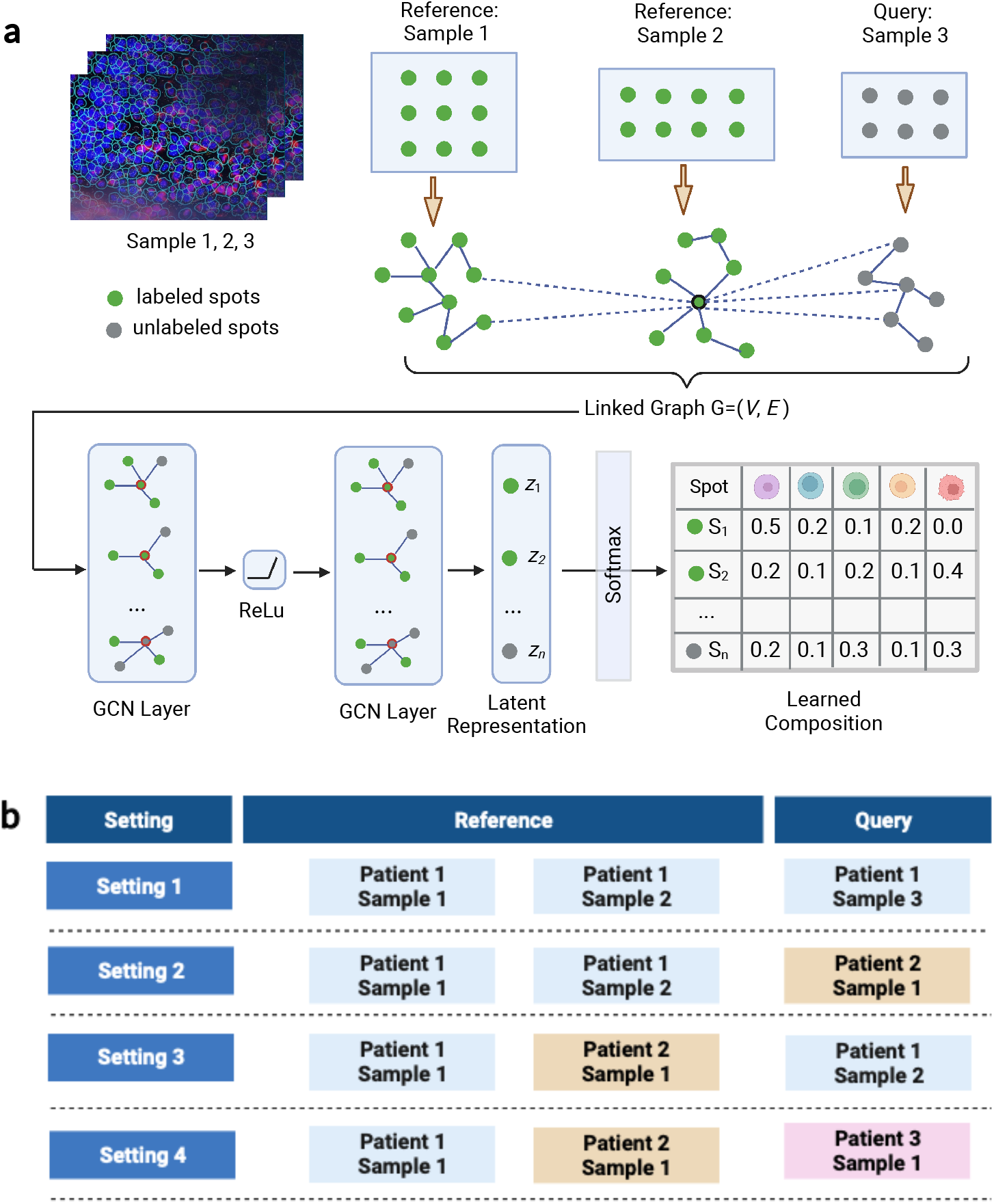
An overview of GNNDeconvolver and experimental settings. **a**, An overview of GNNDeconvolver. Note that reference refers to the training data with known cell type proportion labels, and query indicates test data the model will predict. **b**, Four types of experimental settings.

To leverage spatial information among spots in the reference data with established cell type proportions, we conduct our experiments with training on ST reference samples with known cell compositions and testing on unlabeled query samples. To distinguish the difficulty of the experiments, we divide them into four settings based on the dissimilarity between the reference and query samples, and all of four settings are supported by GNNDeconvolver. Setting 1 only includes samples from the same patient in both reference and query. In setting 2, the reference samples are from the same patient, while the query samples are from other patients. In setting 3, reference samples are from different patients and the sample patients in the query are present in the reference. Unlike setting 3, the query sample patients in setting 4 have no previous reference (Fig. 3).

Based on totally 20 samples in SpatialCTD across three human tissues, we assess the performance of GNNDeconvolver and state-of-the-art methods in the following four perspectives: (i) overall prediction accuracy, which is the average of the measured metrics of Mean Squared Error (MSE), Mean Absolute Error (MAE), and Pearson Correlation Coefficient (PCC); (ii) performance of each individual metric (MSE, MAE and PCC); (iii) performance on each tissue (Lung, Kidney and Liver); and (iv) performance under each setting (Setting 1, 2, 3 and 4); Our results show that GNNDeconvolver often outperforms existing state-of-the-art methods, followed by RCTD and CARD (Fig. 4). A detailed description about how to calculate rankings in terms of each perspective is provided below.

**Figure 4:**
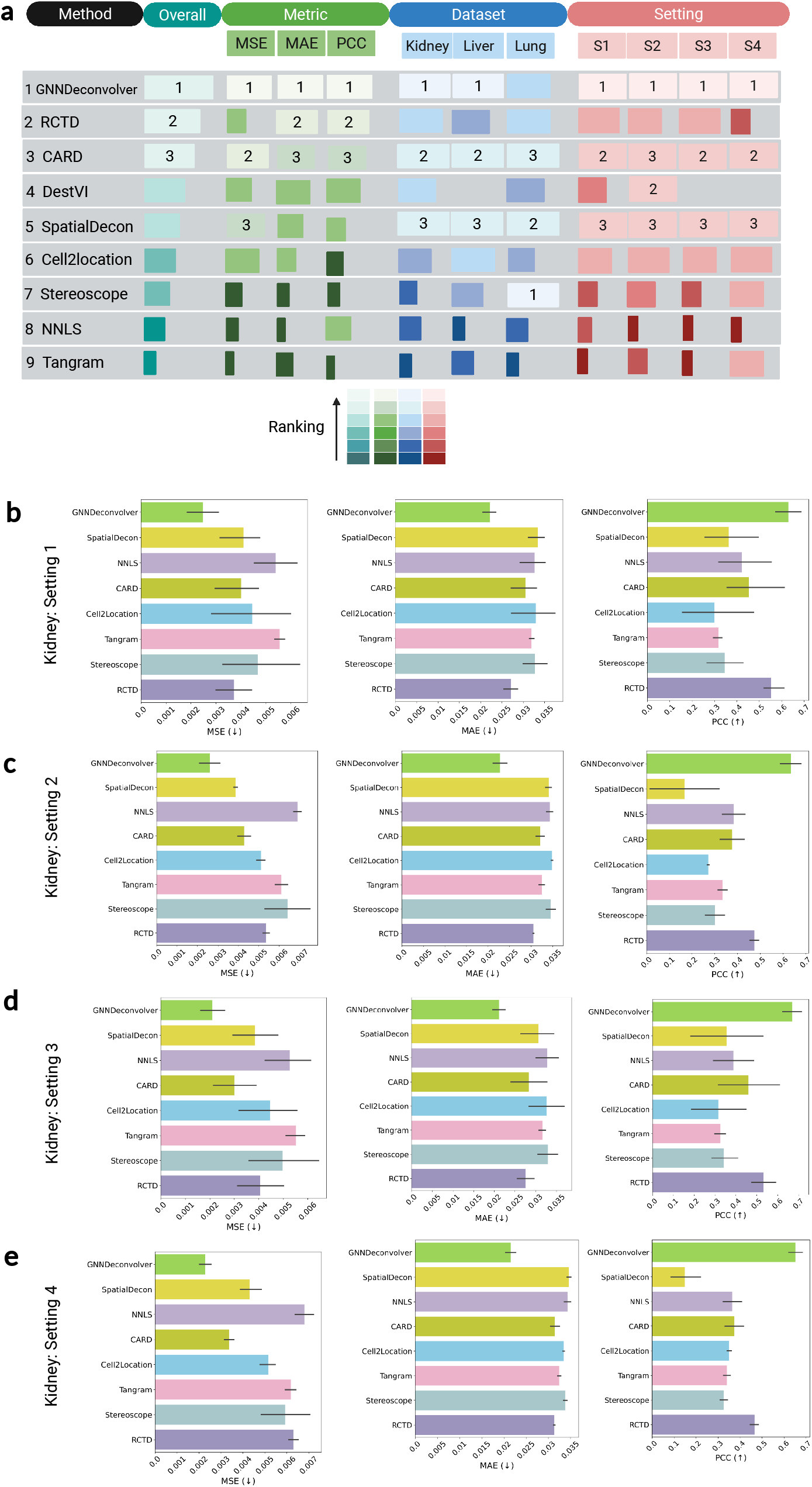
Performance of 9 methods in cell type deconvolution. **a**, A summary of results for 9 cell type deconvolution methods. **b**, Comparison of the models under different settings on SpatialCTD lung tissue in terms of MSE, MAE and PCC.

Step 1: MSE, MAE and PCC are computed for each method across all experimental settings.

Step 2: For each method under each experiment, three ranking scores are established based on its performance in terms of MSE, MAE, and PCC.

Step 3: We then calculate the average metric specific rankings under all experiments to represent this method’s overall metric ranking (ii).

Step 4: The overall ranking for each method is determined by the average ranking of overall MSE, MAE and PCC ranking mentioned above (i).

For overall ranking under each tissue, we would repeat above steps and replace all experiments in Step 3 with tissue specific experiments (iii). Similarly, to calculate overall ranking under each setting, we repeat above steps and replace all experiments in Step 3 with setting specific experiments (iv).

We find that the accuracy of most of the 9 methods is generally stable across three metrics, three tissues and four settings. In terms of overall ranking, GNNDeconvolver (1.57) outperforms existing representative methods by a substantial margin, like RCTD (3.27), CARD (3.28) and DestVI (4.7). GNNDeconvolver performs consistently to achieve the first place among all methods in most cases (MSE:1.65, MAE:1.66, PCC: 1.4, Kidney:1.1, Liver: 1.25, Setting 1:1.8, Setting 2: 2.5, Settig 3: 1.3 and Setting 4: 1.0). CARD is always ranked second or third regardless of which evaluation aspect. However, the accuracy of some methods (i.e. Cell2location or stereoscope) varies depending on tissues and settings (Fig. 4). In conclusion, GNNDeconvolver usually outperforms baselines tremendously and its performance is very robust across various settings. These achievements of GNNDeconvolver can be attributed to its following advantages. First, GNNDeconvolver is robust to noise and dropout in ST data where certain gene expression measurements in spots are missing or unreliable, as GNNDeconvolver can leverage the relationships between spots and their neighbors to better estimate missing or noisy values. Second, GNNDeconvolver has the potential ability of transfer learning. GNNDeconvolver can be trained on reference samples with known cell type composition and applied to other samples, allowing it to learn from multiple sources and adapt to different experimental conditions. This capability enables GNNDeconvolver to leverage prior knowledge in reference samples and generalize across different ST samples. Third, the cell type proportions in neighboring spots exhibit a high degree of similarity. GNNDeconvolver can incorporate the spatial information of spots into graph construction, and can capture local relationships among spots by leveraging the graph structure, which help distinguish cell type proportions among spots.

Specifically, on the kidney tissue, there are 4, 2, 6 and 8 experiments conducted and evaluated in Setting 1, 2, 3 and 4 respectively in terms MSE, MAE and PCC (Fig. 4). For MSE, we find that GNNDeconvolver consistently ranks first while CARD and RCTD are among the top three methods across all settings. GNNDeconvolver consistently achieves the highest MAE and PCC scores, while RCTD consistently ranks second for both MAE and PCC metrics no matter which setting. Simultaneously, we observe that RCTD demonstrates favorable performance in the first three settings (S1, S2 and S3), but exhibits suboptimal performance compared to the majority of the other methods in the last setting (S4) in terms of MSE. This may be attributed its limited generalization ability given that both reference and query samples in S4 are derived from distinct patients. RCTD assumes that the proportion of genes in each spot correlates with the distribution of cell types in that spot, thereby relying on specific data distribution assumptions. Nevertheless, the data distribution across patients may exhibit considerable variability. Our analysis also suggests that the performance of a given method may vary across different metrics. For instance, while RCTD yields satisfactory results in PCC and MAE across all four settings, its performance in MSE is inferior to that in PCC and MAE. GNNDeconvolver can still exhibit exceptional performance on the liver tissue, but its efficacy is reduced when applied to the lung tissue (Fig. 17 in Supplementary).

To further demonstrate the efficacy of GNNDeconvolver, we reconstruct spatial maps of cell type distributions for the SpatialCTD lung 5-2 sample dataset using the deconvolution proportions generated by the model. Subsequently, we compare these maps with the established gold standard. We can clearly see that the spatial patterns of cell types predicted by GNNDeconvolver are consistent with the ground truth (Fig. 5). To be specific, GNNDeconvolver correctly maps the tumor cells (red) at the bottom left and spreading at upper right area and B cells (blue) on the diagonal. What’s more, we explore spatial distribution of cell type proportion from each cell type (Fig. 5). The first striking pattern is the similar spatial organization between macrophage cells and neutrophil cells, which indicates the strong extent of co-localization between two cell types in the whole sample. Similar colocalization pattern can be found between CD4 memory cells and CD8 memory cells. We also find that tumor cells and macrophage cells are arranged mutually exclusive in space. This is consistent with the conclusion from previous studies of tumor-associated macrophages in lung tumor [26]. The study typifies the complexity of macrophage function in cancer which is the respective antitumoral and protumoral roles of M1 and M2 tumor-associated macrophages. In spatial distribution of cell type proportion for macrophage cells, we interpret the dark dots represent M1 macrophage cells playing an antitumoral role while light colored dots indicate M2 macrophage cells playing a protumoral role.

**Figure 5:**
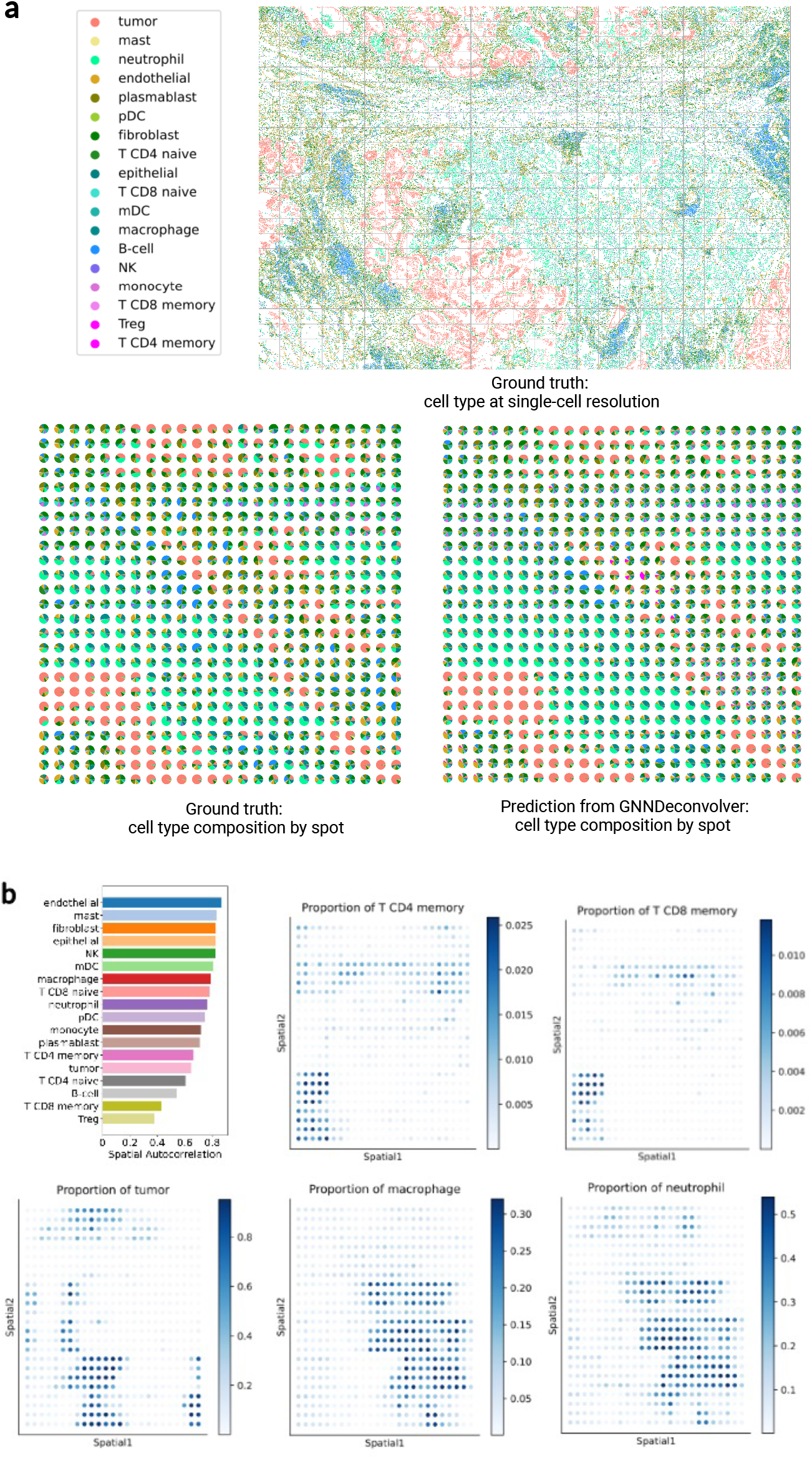
GNNDeconvolver deconvolution on SpatialCTD lung 5-2 sample. **a**, Ground truth singlecell resolution on SpatialCTD Lung 5-2 sample. Each dot is a single cell colored by its ground truth cell type label. Proportions of deconvolved cell types from ground truth and GNNDeconvolver represented as pie charts for each spot. **b**, Spatial autocorrelation of the cell type proportion computed using Hotspot. Spatial distribution of cell type proportion for T CD4 memory cells, T CD8 memory cells, tumor, macrophage and neutrophil cells, as inferred by GNNDeconvolver. Each dot represents a spot. The depth of the point indicates the proportion of the cell type in the spot.

### 2.5 Impact of spot size

To evaluate the stability of the method with respect to spot size of the spatial transcriptomics (ST) data, we test all methods on three different spot sizes (large spots: 32338.2067 *μm*^2^), medium spots: 16169 *μm*^2^ and small spots: 10779 *μm*^2^). We find that GNNDeconvolver consistently performs the best among all different spot sizes no matter which metric evaluation and is more stable to varying spot sizes (Fig. 6). This nice property of GNNDeconvolver is enbaled by its intrinsic design. GNNDeconvolver aggregates information and labels from neighboring nodes within the graph structure, thus it can adapt to different spot sizes through the efficient integration of information from diverse numbers of neighboring nodes. In terms of MSE, the performance of all methods except CARD tends to become increasingly worse when the spot size becomes smaller, which is consistent with the results in previous cell type deconvolution benchmarking studies [25]. This phenomenon can be also observed on the PCC metric for all methods. One difference between these two metrics is that the variation of PCC is not as obvious as that of MSE for different spot sizes. Note that the PCC score of SpatialDecon and RCTD drops greatly when spot size decreases from the large spot size to the medium and small spot sizes. What’s more, different from other methods, both Tangram and RCTD perform best in small spots but not in large spots in terms of the MAE metric. The MAE of all methods performs similarly on different spot sizes, unlike MSE and PCC, which are more sensitive to the spot size.

**Figure 6:**
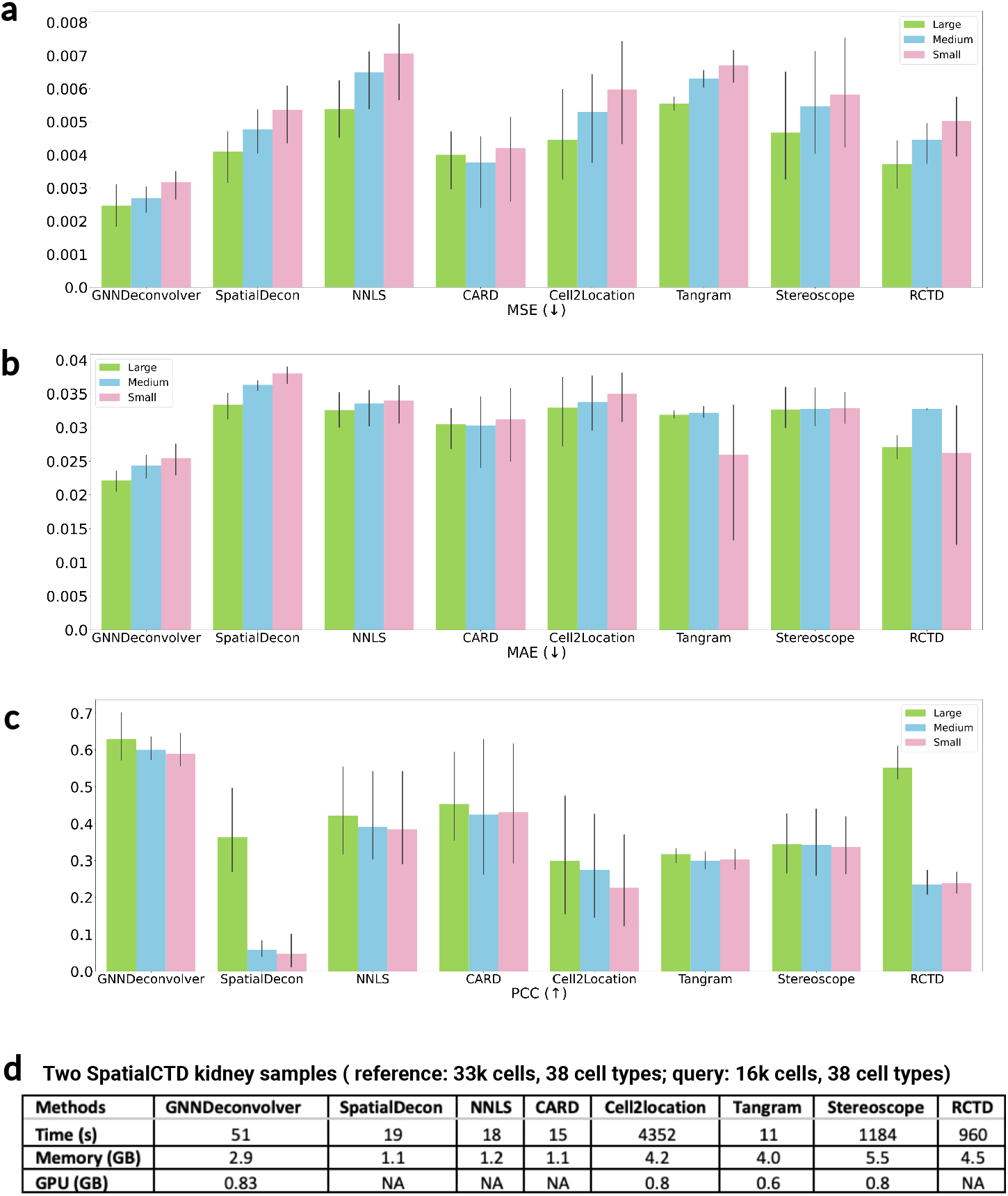
Assessment of 9 methods from stability and usability perspectives. **a-c**, Comparison of 9 models on different pseudo spots size for SpatialCTD kidney Setting 1. The determination of spot size relies on the method of dividing one field of view (FOV). We conduct simulations of three distinct spot sizes, where each FOV is partitioned into 20 (4×5) spots denoted as large spots (spot area: 32338.2067 *μm*^2^), 40 (5×8) spots as medium spots (spot area: 16169 *μm*^2^), and 60 (6×10) spots as small spots (spot area: 10779 *μm*^2^). **d**, Usability evaluation of the methods in terms of running time, system memory and GPU memory. All experiments are conducted in the same platform (12.7 GB System Ram, 166.8 GB Disk, 46080 KB Cache, 6 CPU cores, Tesla T4 GPU).

### 2.6 Computational resources

In order to assess the usability, we conduct comprehensive investigations of all methods in terms of running time, system memory, and GPU memory. The experiment is conducted on SpatialCTD kidney datasets, where the reference data (training data) is the sample Run1087 SP20 1098 with 33k cells, and the query data (test data) is the sample Run1087 SP19 4061 with 16k cells. All experiments are conducted on the same platform (12.7 GB System Ram, 166.8 GB Disk, 46080 KB Cache, 6 CPU cores, Tesla T4 GPU) for fair comparison. It is noteworthy that certain methods, such as GNNDeconvolver and Tangram, rely on deep learning techniques and can potentially benefit from GPU acceleration.

Regarding overall running time, we observe that Tangram has shortest running time, which is consistent with the conclusion from previous cell type deconvolution benchmarking studies [25]. The running time for CRAD, NNLS, SpatialDecon, and GNNDeconvolver is under 60 seconds. Conversely, Cell2location requires around 20 minutes to complete, and Sterescope typically runs more than one hour (Fig. 6). It should be noted that GNNDeconvolver not only demonstrates superior performance relative to all other methods, but also is capable of completing its computations in less than one minute.

Regarding system memory consumption and GPU memory consumption, our findings indicate that Tangram, Stereoscope, and RCTD exhibit higher RAM requirements, whereas SpatialDecon, NNLS, and CRAD exhibit lower RAM requirements. The RAM requirements of GNNDeconvolver fall within an intermediate range. GNNDeconvolver, Cell2location, Tangram and Stereoscope have similar GPU memory consumption while others only work under CPU computation.

In summary, CARD exhibits the highest level of efficiency, whereas Stereoscope and Cell2location are the least efficient. Among the top-performing methods, GNNDeconvolver and CARD are the most efficient.

## 3 Discussion

The advent of emerging spatial transcriptomics (ST) technologies has provided novel insights into cellular abundance and gene expression heterogeneity in a spatial context. Nevertheless, the resolutions of most of the current ST data are not guaranteed to be single-cell. The task of cell type deconvolution has been developed to isolate discrete cell types from such multicellular spots. However, the existing datasets for cell type deconvolution are either simulated from single-cell RNA seq data lack of biological variance or obtained from ST data at a single-cell resolution but constrained in scale, primarily comprising data on mice and not intended for human immuno-oncology. Therefore, we introduce a comprehensive spatial transcriptomic dataset called SpatialCTD, comprising 1.8 million cells derived from the human tumor microenvironment in the lung, kidney, and liver. In addition, we introduce a robust Graph Neural Network (GNN)-based approach, known as GNNDeconvolver, that often surpasses current state-of-the-art methods by a significant margin, even in the absence of single-cell RNA-seq data. To facilitate cell type deconvolution evaluations, we provide an online tool capable of converting spatial transcriptomic data at the single-cell resolution into pseudo spots with adjustable spot size.

To demonstrate its necessity, we compare, through both biological variance and statistical differences, SpatialCTD with existing simulated cell type deconvolution benchmark datasets. The results of the differential expression (DE) analysis indicate that some cell types exhibit disparate expression patterns between the natural tumor environment (SpatialCTD data) and the healthy tissue environment (simulated benchmark datasets). We then conduct a thorough comparison of SpatialCTD with a number of existing cell type deconvolution benchmark datasets. Most of the ST data generated via the grid spot way are solely on mice and do not involve human samples while other datasets are simulated from single-cell RNA sequencing data rather than actual ST data, and are not derived from TME tissues neither. It is worth noting that the datasets under comparison have a limited sample size, with the largest comprising only 60,000 cells, whereas SpatialCTD comprises up to 770,000 cells within a single tissue, and sum up to 1.8 millions of cells. What’s more, SpatialCTD presents several distinctive features that are not yet commonly available in existing benchmark datasets above, including spot images, and spatial location data for both cellular and subcellular entities. To ease the process of converting spatial transcriptomics data at single-cell resolution to pseudo spots for cell type deconvolution studies, we have developed a web-based online conversion tool.

In addition, we propose GNNDeconvolver as a Graph Neural Network-based approach that serves as a reliable and strong baseline method for analyzing SpatialCTD. Kindly note that GNNDeconvolver is a method that does not require single-cell RNA seq data as reference data. Extensive experiments show that GNNDeconvolver usually outperforms state-of-the-art methods by a large margin.

To evaluate the stability of the method with respect to spot size of the ST data, we evaluate all methods on three different spot sizes (large spots: 32338.2067 *μm*^2^), medium spots: 16169 *μm*^2^ and small spots: 10779 *μm*^2^). We observe that GNNDeconvolver consistently outperforms all baselines across various spot sizes under each evaluation metric. We also test the running time, system memory and GPU memory usage of all methods in order to assess their usability. We find that GNNDeconvolver and CARD are the most efficient methods among the top-performing methods.

Subsequent studies may incorporate single-cell resolution ST data from other platforms into SpatialCTD. Moreover, more human tissues may be included in SpatialCTD to establish it as a benchmark dataset at the atlas level. For GNNDeconvolver, approaches to combine single-cell RNA seq data and ST data to infer cell type deconvolution can be further explored.

## 4 Methods

### 4.1 SpatialCTD generation

#### Dataset collection

We collect spatial transcriptomic data at a single-cell resolution and subcellular (RNA) resolution from CosMx platform [10] using spatial molecular imaging technique. Twenty samples across human lung, kidney and liver tissues containing tumor cells are obtained.

##### Human Lung

This dataset is generated with a 960-plex CosMx RNA panel run on CosMx SMI. There are 8 samples from 5 NSCLC (non-small cell lung cancer) tissues. It includes 3 female and 2 male patients, all of whom are white and over the age of 60. In total, there are 800,327 cells in this dataset, 766,313 cells are analyzed and 259,604,214 transcripts are detected. Each cell is represented by 960 selected gene targets, and each cell in the tissue is categorized as one of 18 distinct cell types.

##### Human Kidney

This dataset is generated with 960-plex CosMx RNA panel run on CosMx SMI, and is collected from a kidney core biopsy from lupus nephritis patients. There are 10 samples and around 300,000 single cells in this dataset.

##### Human Liver

The CosMx SMI human liver dataset offers a subcellular expression map of 1,000 genes. With over 800,000 single cells and approximately 700 million transcripts, this comprehensive dataset covers a 180 *mm*^2^ area of liver tissue. This dataset contains one sample each from normal liver and hepatocellular carcinoma tissues. Normal liver is from a male patient at the age of 35 while Hepatocellular carcinoma is from a female patient at the age of 65.

#### Generation of pseudo spots

In order to obtain multi-cell-per-spot datasets with known cell type compositions at each spot, we grid our collected single-cell resolution spatial transcriptomics datasets to generate pseudo spots. We aggregate the expression profiles of all cells in the pseudo spot to denote the expression profile of a simulated spot, and sum cell types of all cells residing in the simulated spot to be cell type propositions of the spot as the ground truth. The centered point of the grid spot is defined as the coordinate of the spot. Spots with few cells will be filtered out. After generation, each sample comes with at least five files. Spot cell state file summarizes the number of cells in each pseudo spot. Cell Id mapping file indicates the mapping relationship between cell id and pseudo spot id. Spot location file tells the spatial information of each generated pseudo spot. Spot gene expression file is the spot level gene expression while the ground truth file describes the cell type compositions for each spot. For spot size, we refer to the spot size from GeoMx platform whose mean spot size is 37456.28 *μm*^2^, and median spot size is 24168.74 *μm*^2^.

##### Human Lung

The sizes of all FOVs in human lung samples are identical (5,472 pixels * 3,648 pixels, 0.18 *μm* per pixel). We divide each FOV into 20 (4*5) pseudo spots. Each spot area is 32338.2067 *μm*^2^ within the reasonable range of GeoMx spot size. After filtering out generated spots with low quality, we totally get 4,660 generated spots from 8 samples and 771,236 cells are kept.

##### Human Kidney

The sizes of all FOVs in human kidney samples are identical (5,472 pixels * 3,648 pixels, 0.18 *μm* per pixel). Same with human lung, we divide each FOV into 20 (4*5) pseudo spots. Each spot area is 32338.2067 *μm*^2^. Once spots with low quality have been filtered out, we totally get 2,460 spots from 10 samples, and 296,838 cells are kept.

##### Human Liver

The sizes of all FOVs in human liver samples are identical (4,236 pixels * 4,236 pixels, 0.12 *μm* per pixel). Each FOV would be divided into 9 (3*3) pseudo spots. Each spot area is 28709.9136 *μm*^2^. After filtering out generated spots with low quality, we totally get 5,796 generated spots from 2 samples and 760,506 cells are kept.

### 4.2 GNNDeconvolver

Assuming that we have *t* samples with *n*_1_, *n*_2_,…, *n*_*t*_ spots respectively, and the gene expression dimension of each spot is identical as *d*. We consider each spot as a node in a graph. For each sample, we construct individual graph connecting all nodes in the sample via spatial distance and spot gene expression. To be specific, we first construct two adjacency matrices *A*_spatial_ and *A*_gene_ based on spatial distance and gene expression, respectively. For *A*_spatial_, each node is connected to its closest top *K* nodes in the surrounding space. Here we can also set the threshold to connect pair of nodes. When the distance between two nodes is less than the specified threshold, the two nodes would be connected. Similarly, *A*_gene_ is built up by connecting node with its most similar top *K* nodes of gene expression. Then the final adjacency matrix *A*_sample_ of each sample is obtained by weighting two adjacency matrices *A*_sample_ = *αA*_spatial_ + *βA*_gene_ where *α* and *β* are weights. The graph for sample 1 can be represented as 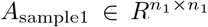 and 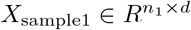 where *A*_sample1_ is the final adjacency matrix, and *X*_sample1_ is the node representation matrix.

To connect nodes across samples, we only leverage gene expression to construct the KNN graph since spatial information of spots across samples doesn’t exist. To be specific, to connect nodes in sample 1 with nodes in other samples, we first calculate the gene expression similarity between the node in sample 1 and all nodes in other samples. We then choose the *K* most similar nodes across samples and establish connections between them. In such case, we can construct one graph connecting nodes across all *t* samples where some samples are labeled with cell type compositions and some are not. The graph can be represented as *A*_all_ ∈ *R*^*n×n*^ and *X*_all_ ∈ *R*^*n×d*^ where *n* = *n*_1_ + *n*_2_ + ⋯ + *n*_*t*_. Next, we will introduce how our designed Graph Neural Network based method (GNNDeconvolver) propagates features from labeled nodes to infer cell type compositions of unlabeled nodes in a graph.

After graph construction, we have a linked graph *G* = (*V, E*). The goal of GNNDeconvolver is to predict cell type proportions of unlabeled spots by using not only the features of each spot, but also the graph structure describing the connections between labeled nodes and unlabeled nodes, which is characterized as the above adjacent matrix *A*. To be clear and specific, GNNDeconvolver takes two inputs. One is the adjacent matrix *A* ∈ *R*^*n×n*^ learned from the graph construction process. The other is the node representation matrix *X* ∈ *R*^*n×d*^ where *n* indicates the number of spots and *d* is the number of variable genes.

GNNDeconvolver consists of two convolutional layers. Specifically, each graph convolutional layer is defined as:

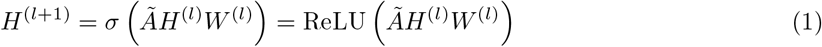

where 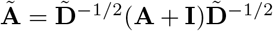 with 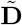 the diagonal matrix of **A** + **I** and **I** the identity matrix. *H*^(*l*)^ is the input from the previous layer. *W* ^(*l*)^ is the weight matrix of the *l*-th layer. ReLU(·) is the nonlinear activation function. Here, the input for the first layer would be the original node representation H^(0)^ = X.

The overall structure of GNNDeconvolver is described as:

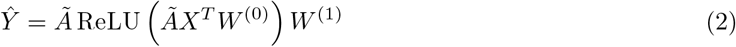

where *W* ^(0)^ and *W* ^(1)^ are weight matrices to be learned, and *Ŷ* is the predicted cell type compositions with *F* unique cell types. The loss function is defined as the cross-entropy at labeled spots below:

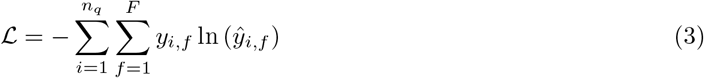

where *n*_*q*_ is the number of all labeled nodes from samples, *ŷ*_*i,f*_ represents the predicted value of cell type *f* in spot *i*, and *y*_*i,f*_ indicates real proportion value of cell type *f* in spot *i*. The model is trained by minimizing the cross-entropy *L* between the known cell type compositions and the predicted cell type proportions on training datasets, and parameters are learned via stochastic gradient descent.

### 4.3 Baselines

#### SpatialDecon

We used a python implementation https://github.com/OmicsML/dance/blob/main/dance/modules/spatial/cell_type_deconvo/spatialdecon.py that follows the guidelines on the SpatialDecon (R) GitHub repository: https://github.com/Nanostring-Biostats/SpatialDecon.

#### CARD

We used a python implementation https://github.com/OmicsML/dance/blob/main/dance/modules/spatial/cell_type_deconvo/card.py that follows the guidelines on the CARD (R) GitHub repository: https://github.com/YingMa0107/CARD. We applied hyper-parameter tuning to optimize this model, which has default parameters max_iter=100, converge threshold epsilon=1e-4, and spatial gaussian scale factor sigma=1e-1.

#### RCTD

We followed the guidelines on the RCTD GitHub repository: https://github.com/dmcable/RCTD. We set doublet_mode = ‘full’.

#### Cell2location

We followed the guidelines on the Cell2location GitHub repository: https://github.com/BayraktarLab/cell2location. The single-cell model was trained with parameters max epochs = 250 and lr = 0.002. The Cell2location model was obtained with parameters max_epochs = 30,000, N_cells_per_location = 120, and detection_alpha = 20.

#### DestVI

We followed the guidelines on the DestVI GitHub repository: https://github.com/scverse/scvi-tutorials/blob/main/DestVI_tutorial.ipynb. The single-cell model was trained with parameters max epochs = 250, lr = 0.001. The spatial model was trained with max_epochs = 2,500.

#### Stereoscope

We followed the guidelines on the stereoscope GitHub repository: https://github.com/almaan/stereoscope. The single-cell model was trained with parameters max epochs = 100. The spatial model was trained with max epochs = 10,000.

#### NNLS

It is a linear regression model. We followed the guidelines from sklearn: https://scikit-learn.org/stable/modules/generated/sklearn.linear_model.LinearRegression.html. We applied a non-negative regression model, without bias.

#### Tangram

We followed the instructions on the Tangram GitHub repository: https://github.com/bro_adinstitute/Tangram. We set the parameters as modes = ‘clusters’ and density = ‘rna_count_based’. To deconvolute the cell types in space, we invoke ‘project_cell_annotation’ to transfer the annotation to space.

### 4.4 Evaluation metrics

We use the following metrics to evaluate the predicted cell type composition for cell type deconvolution task.

#### MSE

The Mean Squared Error (MSE) describes the average squared difference between the predicted values and the actual values in a dataset. A lower MSE value indicates better prediction accuracy. It is defined as:

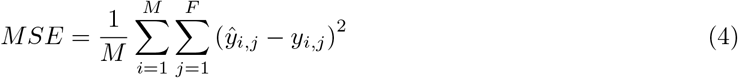

where *M* is the number of all unlabeled spots in the test dataset, and *F* indicates the number of unique cell types to be predicted. *ŷ*_*i,j*_ is the predicted value of cell type *j* in spot *i* while *y*_*i,j*_ is the ground truth.

#### MAE

The Mean Absolute Error (MAE) measures the average magnitude of the errors in a set of predictions. It’s the average over the test sample of the absolute differences between prediction and actual observations where all individual differences have equal weight. A lower MAE value indicates better prediction accuracy. It is defined as follows:

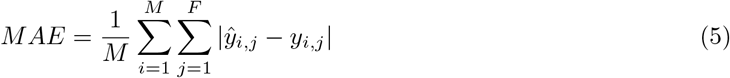

#### PCC

The Pearson Correlation Coefficient (PCC) is a measure of linear correlation between two sets of data. A higher PCC value indicates better prediction accuracy. It’s defined as:

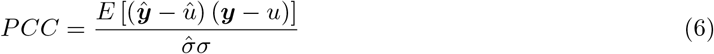

where ***ŷ*** is the predicted cell type proportions with *F* values, and *û* is the average proportion value of the predicted cell types. ***y*** is the ground truth, and *u* is the average proportion value in the ground truth. 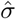 and *σ* are the s.d. of the cell type proportion in the predicted result and ground truth, respectively.

### 4.5 Experimental settings

To incorporate spatial information of spots from reference samples with established cell type proportions, we perform our experiments by training on spatial transcriptomics reference samples with known cell compositions and evaluating the model on unlabeled query samples. To comprehensively evaluate deconvolution methods, we partition our experiments into four distinct settings based on the dissimilarity between the reference and query samples. Setting 1 comprises reference and query samples from the same patient. In setting 2, the reference samples are derived from the same patient, whereas the query samples originate from different patients with reference samples. In setting 3, reference samples are obtained from distinct patients, but the sample patients in the query are present in the reference. In contrast to setting 3, the query sample patient in setting 4 has no prior reference. In SpatialCTD lung, we conduct 3 experiments in Setting 1 and 2 experiments in Setting 2. In SpatialCTD kidney, we conduct 4 experiments in Setting 1, 2 experiments in Setting 2, 6 experiments in Setting 3 and 8 experiments in Setting 4. In SpatialCTD liver, we conduct 8 experiments in Setting 1.

## References

[1] Visium spatial gene expression (10x genomics, 2020).

[2] Hek293t and ccrf-cem cell line mixture data. https://www.ncbi.nlm.nih.gov/geo/query/acc.cgi?acc=GSE174746,.

[3] Human pdac data. https://www.ncbi.nlm.nih.gov/geo/query/acc.cgi?acc=GSE111672,.

[4] Mouse posterior brain 10x visium data. https://support.10xgenomics.com/spatial-gene-exp ression/datasets/1.0.0/V1_Mouse_Brain_Sagittal_Posterior,.

[5] Alma Andersson, Joseph Bergenstråhle, Michaela Asp, Ludvig Bergenstråhle, Aleksandra Jurek José Fernández Navarro, and Joakim Lundeberg. Single-cell and spatial transcriptomics enables probabilistic inference of cell type topography. Communications biology, 3(1):565, 2020.

[6] Tommaso Biancalani, Gabriele Scalia, Lorenzo Buffoni, Raghav Avasthi, Ziqing Lu, Aman Sanger, Neriman Tokcan, Charles R Vanderburg, °Asa Segerstolpe, Meng Zhang, et al. Deep learning and alignment of spatially resolved single-cell transcriptomes with tangram. Nature methods, 18(11): 1352–1362, 2021.

[7] Ruben Dries, Qian Zhu, Rui Dong, Chee-Huat Linus Eng, Huipeng Li, Kan Liu, Yuntian Fu, Tianxiao Zhao, Arpan Sarkar, Feng Bao, et al. Giotto: a toolbox for integrative analysis and visualization of spatial expression data. Genome biology, 22:1–31, 2021.

[8] Marc Elosua-Bayes, Paula Nieto, Elisabetta Mereu, Ivo Gut, and Holger Heyn. Spotlight: seeded nmf regression to deconvolute spatial transcriptomics spots with single-cell transcriptomes. Nucleic acids research, 49(9):e50–e50, 2021.

[9] Zhiwei Fan, Yangyang Luo, Huifen Lu, Tiangang Wang, YuZhou Feng, Weiling Zhao, Pora Kim, and Xiaobo Zhou. Spascer: spatial transcriptomics annotation at single-cell resolution. Nucleic Acids Research, 51(D1):D1138–D1149, 2023.

[10] Shanshan He, Ruchir Bhatt, Brian Birditt, Carl Brown, Emily Brown, Kan Chantranuvatana, Patrick Danaher, Dwayne Dunaway, Brian Filanoski, Ryan G Garrison, et al. High-plex multiomic analysis in ffpe tissue at single-cellular and subcellular resolution by spatial molecular imaging. bioRxiv, pages 2021–11, 2021.

[11] Thomas N Kipf and Max Welling. Semi-supervised classification with graph convolutional networks. arXiv preprint arXiv:1609.02907, 2016.

[12] Vitalii Kleshchevnikov, Artem Shmatko, Emma Dann, Alexander Aivazidis, Hamish W King, Tong Li, Rasa Elmentaite, Artem Lomakin, Veronika Kedlian, Adam Gayoso, et al. Cell2location maps fine-grained cell types in spatial transcriptomics. Nature biotechnology, 40(5):661–671, 2022.

[13] Bin Li, Wen Zhang, Chuang Guo, Hao Xu, Longfei Li, Minghao Fang, Yinlei Hu, Xinye Zhang, Xinfeng Yao, Meifang Tang, et al. Benchmarking spatial and single-cell transcriptomics integration methods for transcript distribution prediction and cell type deconvolution. Nature methods, 19(6): 662–670, 2022.

[14] Anjali Rao, Dalia Barkley, Gustavo S Franca, and Itai Yanai. Exploring tissue architecture using spatial transcriptomics. Nature, 596(7871):211–220, 2021.

[15] Micha Sam Brickman Raredon, Junchen Yang, Neeharika Kothapalli, Wesley Lewis, Naftali Kaminski, Laura E Niklason, and Yuval Kluger. Comprehensive visualization of cell–cell interactions in single-cell and spatial transcriptomics with niches. Bioinformatics, 39(1):btac775, 2023.

[16] Matthew E Ritchie, Belinda Phipson, D. Wu Yifang Hu, Charity W Law, Wei Shi, and Gordon K Smyth. limma powers differential expression analyses for rna-sequencing and microarray studies. Nucleic acids research, 43(7):e47–e47, 2015.

[17] Samuel G Rodriques, Robert R Stickels, Aleksandrina Goeva, Carly A Martin, Evan Murray, Charles R Vanderburg, Joshua Welch, Linlin M Chen, Fei Chen, and Evan Z Macosko. Slide-seq: A scalable technology for measuring genome-wide expression at high spatial resolution. Science, 363 (6434):1463–1467, 2019.

[18] Qianqian Song and Jing Su. Dstg: deconvoluting spatial transcriptomics data through graph-based artificial intelligence. Briefings in Bioinformatics, 22(5):bbaa414, 2021.

[19] Patrik L Ståhl, Fredrik Salmén, Sanja Vickovic, Anna Lundmark José Fernández Navarro, Jens Magnusson, Stefania Giacomello, Michaela Asp, Jakub O Westholm, Mikael Huss, et al. Visualization and analysis of gene expression in tissue sections by spatial transcriptomics. Science, 353 (6294):78–82, 2016.

[20] Robert R Stickels, Evan Murray, Pawan Kumar, Jilong Li, Jamie L Marshall, Daniela J Di Bella, Paola Arlotta, Evan Z Macosko, and Fei Chen. Highly sensitive spatial transcriptomics at near-cellular resolution with slide-seqv2. Nature biotechnology, 39(3):313–319, 2021.

[21] Tim Stuart, Andrew Butler, Paul Hoffman, Christoph Hafemeister, Efthymia Papalexi, William M Mauck, Yuhan Hao, Marlon Stoeckius, Peter Smibert, and Rahul Satija. Comprehensive integration of single-cell data. Cell, 177(7):1888–1902, 2019.

[22] Gregor Sturm, Francesca Finotello, Florent Petitprez, Jitao David Zhang, Jan Baumbach, Wolf H Fridman, Markus List, and Tatsiana Aneichyk. Comprehensive evaluation of transcriptome-based cell-type quantification methods for immuno-oncology. Bioinformatics, 35(14):i436–i445, 2019.

[23] Luyi Tian, Fei Chen, and Evan Z Macosko. The expanding vistas of spatial transcriptomics. Nature Biotechnology, pages 1–10, 2022.

[24] Cameron G Williams, Hyun Jae Lee, Takahiro Asatsuma, Roser Vento-Tormo, and Ashraful Haque. An introduction to spatial transcriptomics for biomedical research. Genome Medicine, 14(1):1–18, 2022.

[25] Lulu Yan and Xiaoqiang Sun. Benchmarking and integration of methods for deconvoluting spatial transcriptomic data. Bioinformatics, 39(1):btac805, 2023.

[26] Xiang Zheng, Andreas Weigert, Simone Reu, Stefan Guenther, Siavash Mansouri, Birgit Bassaly, Stefan Gattenlöhner, Friedrich Grimminger, Soni Savai Pullamsetti, Werner Seeger, et al. Spatial density and distribution of tumor-associated macrophages predict survival in non–small cell lung carcinomatumor-associated macrophage subtypes in lung cancer. Cancer research, 80(20):4414– 4425, 2020.

