## Supplementary file for "SpatialCTD: a large-scale TME spatial transcriptomic dataset to evaluate cell type deconvolution for immuno-oncology"

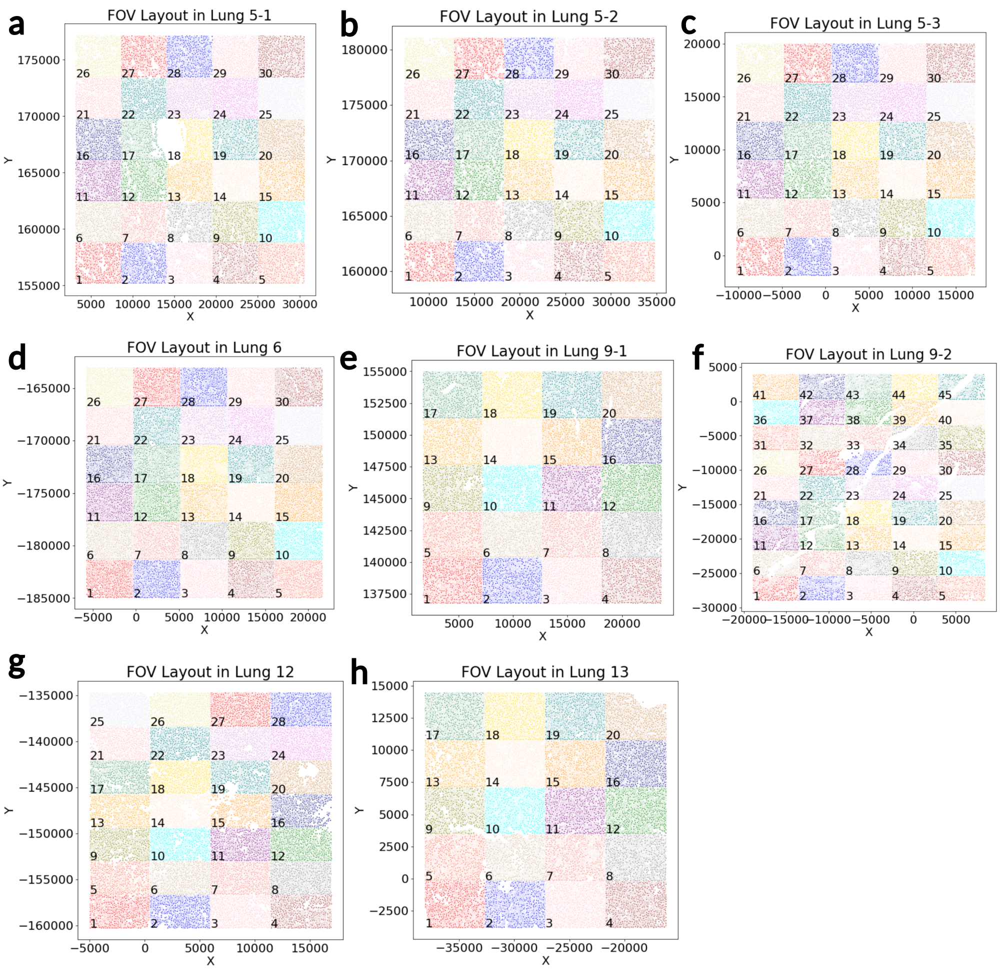


Figure 7: FOV layout of all samples in SpatialCTD human lung tumor.

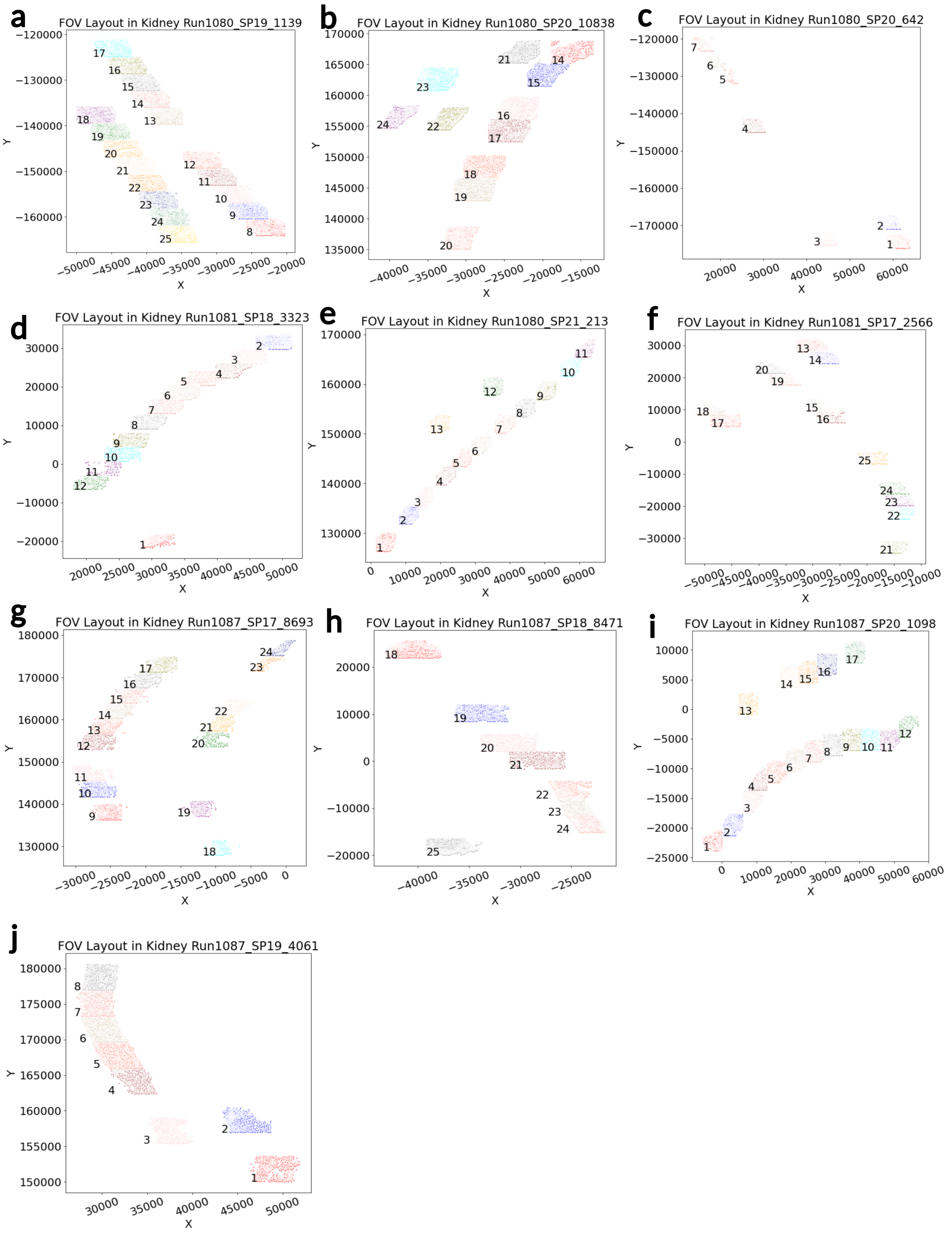


Figure 8: FOV layout of all samples in SPATIALCTD human kidney tumor.



Draft

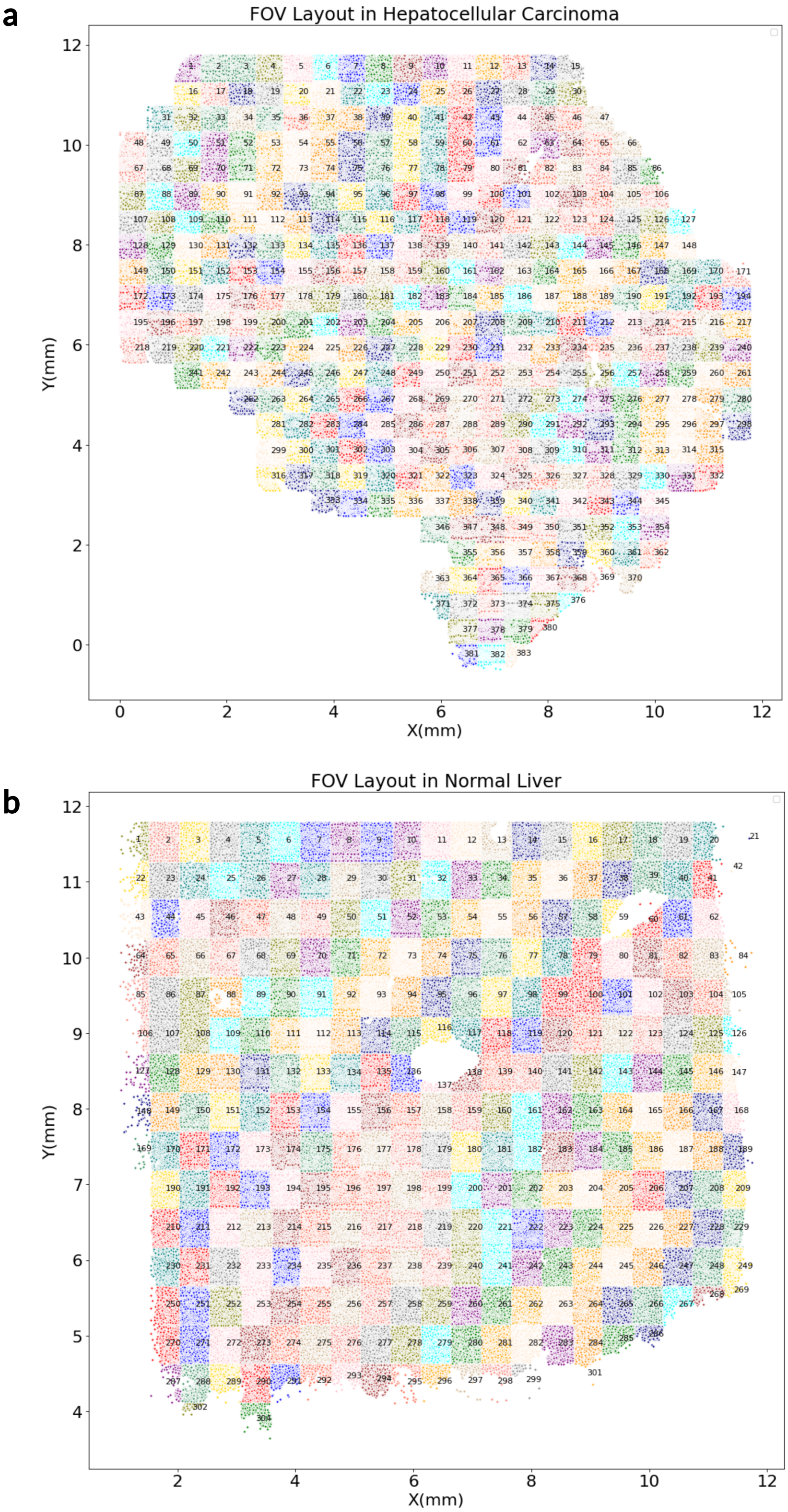


Figure 9: FOV layout of all samples in SpatialCTD human liver.



Draft

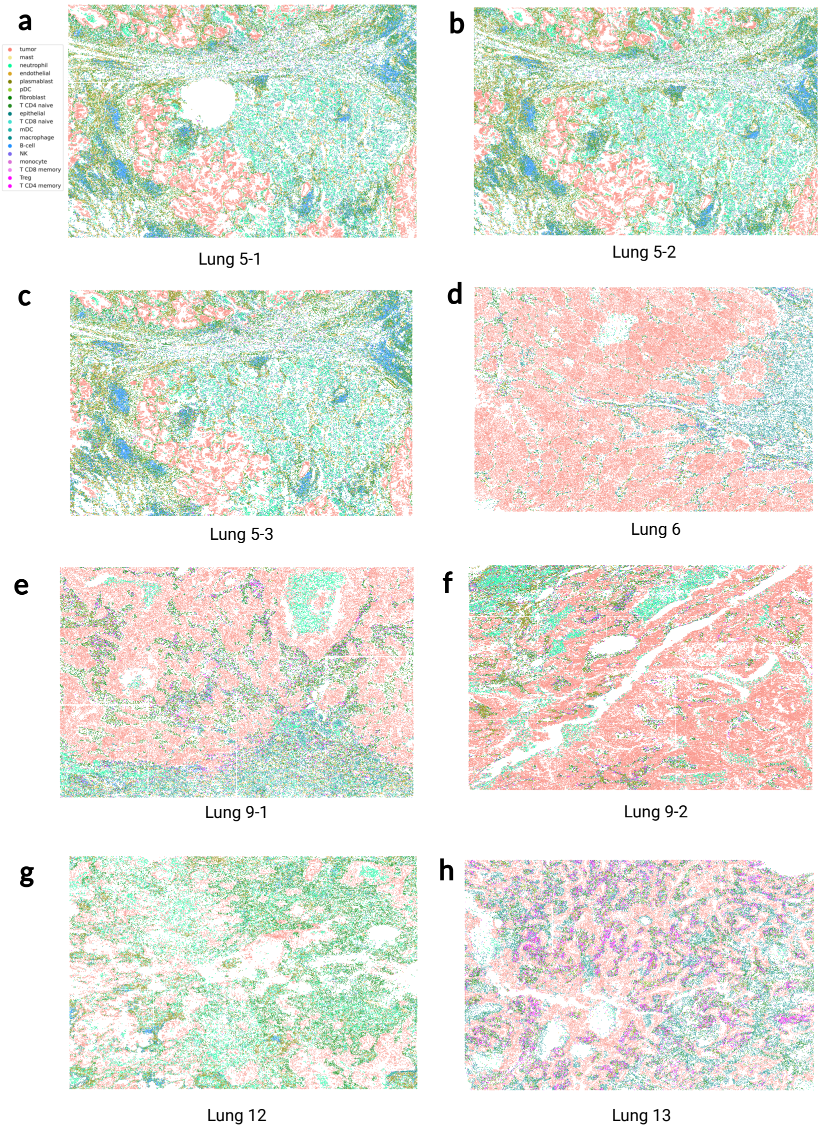


Figure 10: Cell type distribution of each sample in SpatialCTD lung. **a-h**, Ground truth single-cell resolution of all 8 lung samples in SpatialCTD. Each dot is a single cell colored by its ground truth cell type label.

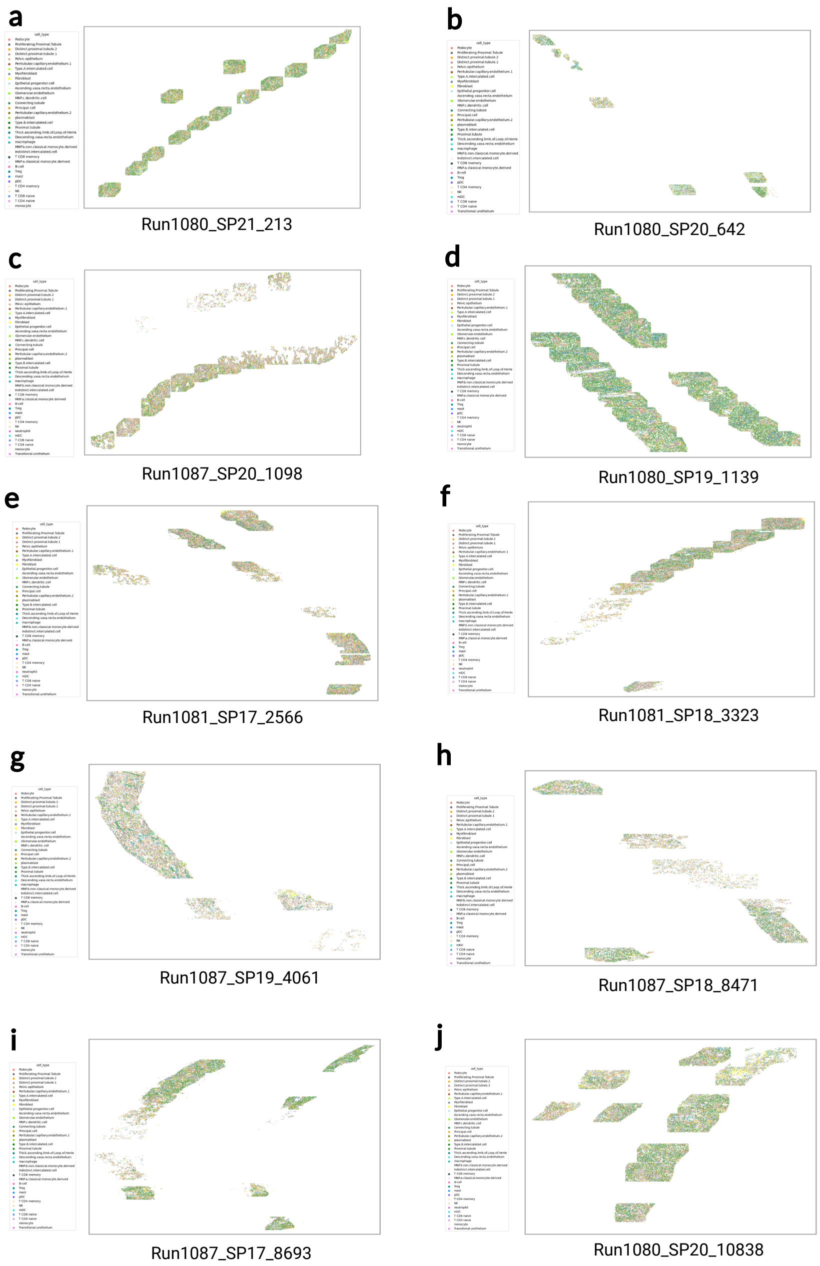


Figure 11: Cell type distribution of each sample in SpatialCTD kidney. **a-j**, Ground truth single-cell resolution of all 10 kidney samples in SpatialCTD. Each dot is a single cell colored by its ground truth cell type label.

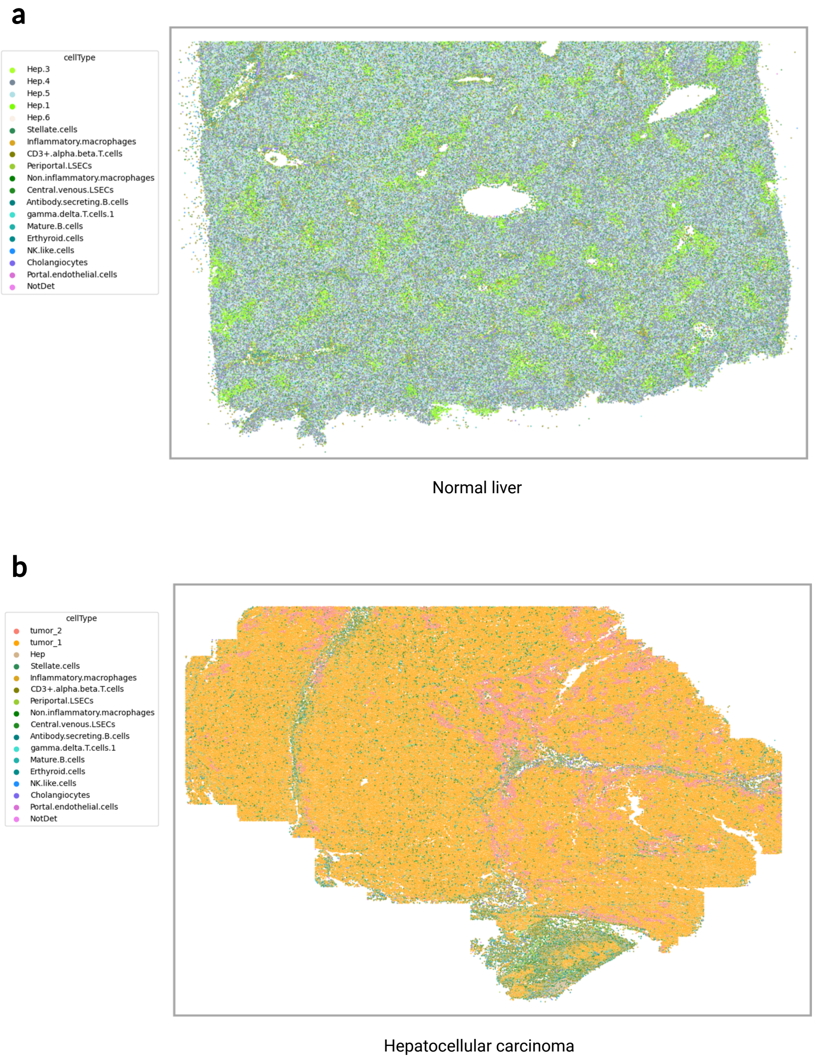


Figure 12: Cell type distribution of each liver sample in SpatialCTD. **a**, Ground truth single-cell resolution of normal liver sample in SpatialCTD. Each dot is a single cell colored by its ground truth cell type label. **b**, Ground truth single-cell resolution of hepatocellular carcinoma sample in SpatialCTD. Each dot is a single cell colored by its ground truth cell type label.

a

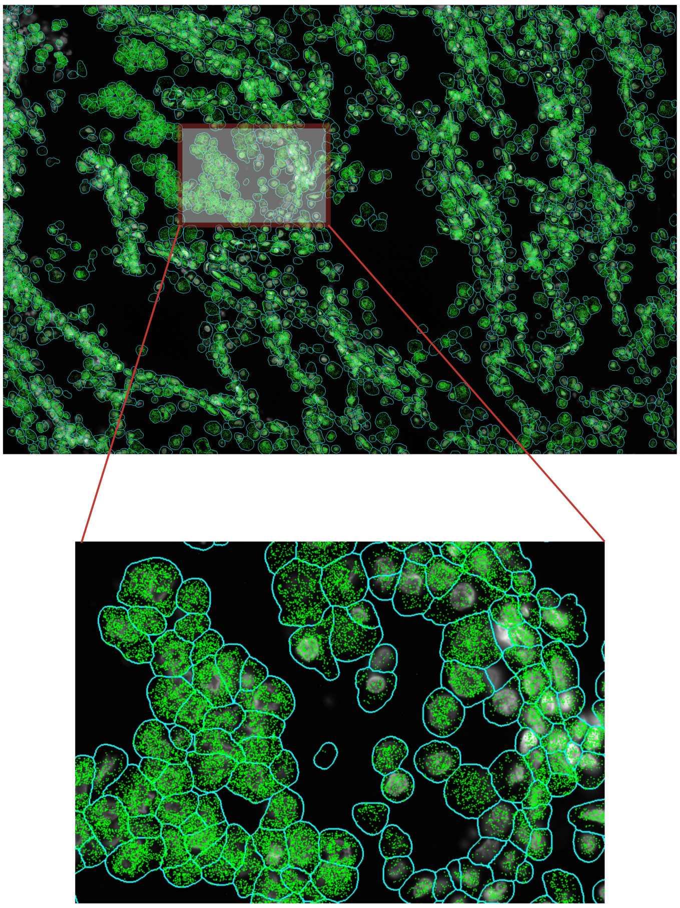

Figure 13: Subcellular spatial transcriptomes in SpatialCTD. **a**, Cell boundaries (cyan) and subcellular spatial transcriptomes represented as dots (green) in cell boundaries, lung 5-1 FOV 1. The picture at the bottom is a zoom in level.



Draft

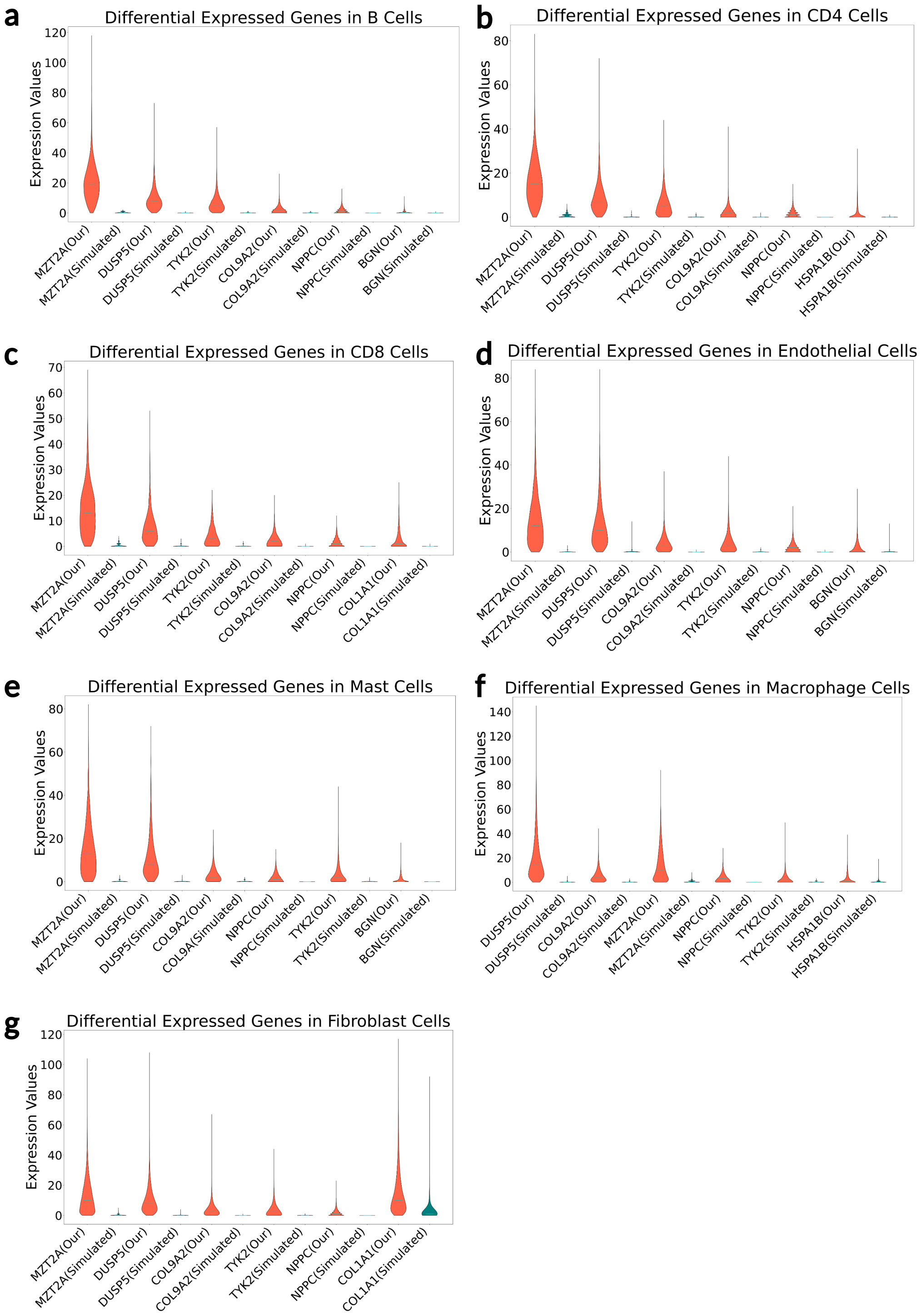


Figure 14: Cellular heterogeneity between actual spatial transcriptomic data (SpatialCTD) and the existing benchmark dataset which is simulated from single-cell RNA sequencing data. **a-g**, A portion of DE genes in macrophage cells, fibroblast cells, B cells, CD4 cells, CD8 cells, endothelial cells and mast cells between SpatialCTD human Lung 5-1 sample and a simulated benchmark dataset from healthy human lung tissue. These genes are highly expressed in our SpatialCTD dataset, but low expressed in the simulated dataset.

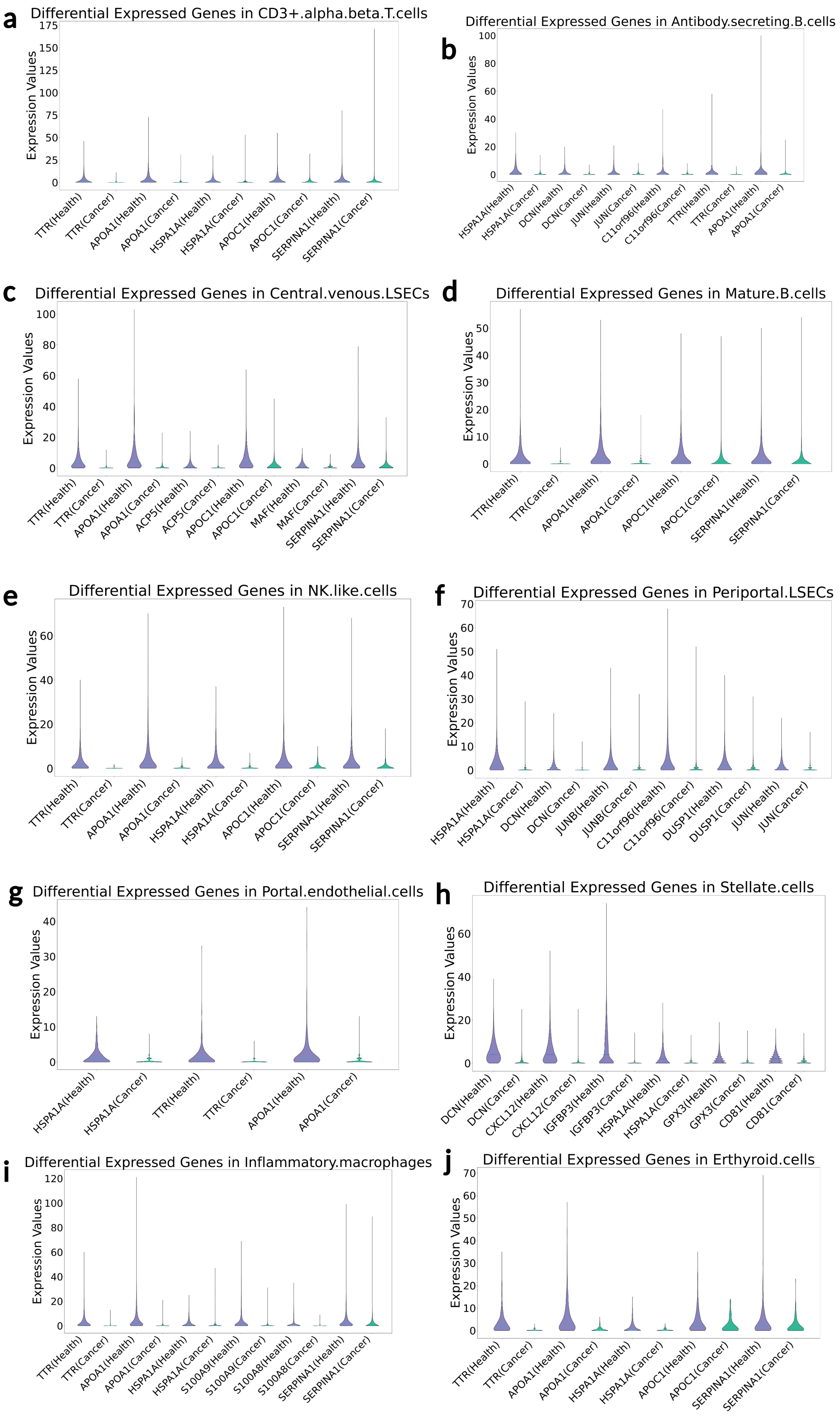


Figure 15: Cellular heterogeneity in SpatialCTD between normal and tumor tissues. **a-j**, A portion of DE genes in several main cell types between SpatialCTD healthy liver tissue sample and hepatocellular carcinoma sample. These genes are highly expressed in SpatialCTD healthy liver tissue, but low expressed in SpatialCTD hepatocellular carcinoma.

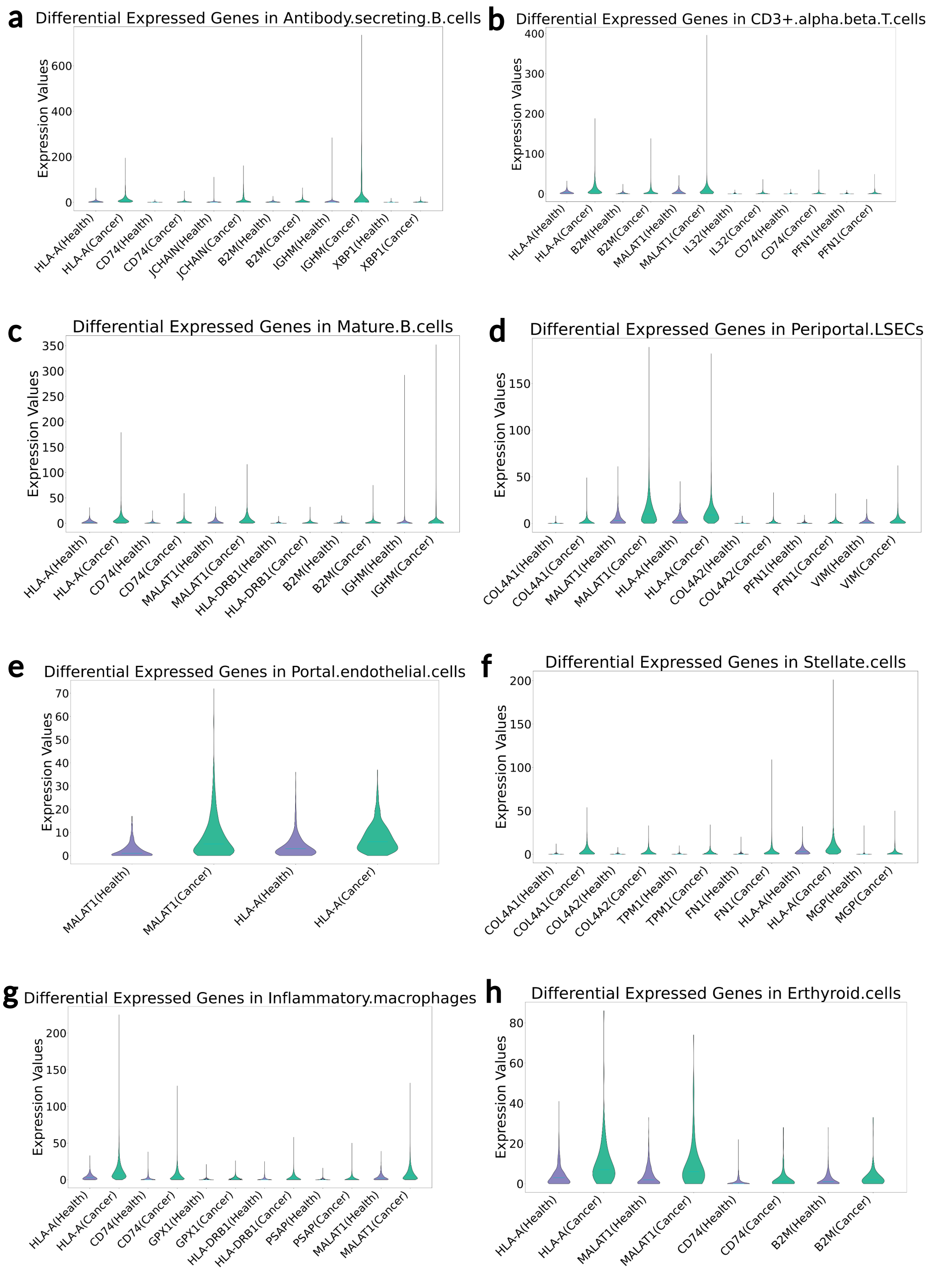


Figure 16: Cellular heterogeneity in SpatialCTD between normal and tumor tissues. **a-h**, A portion of DE genes in several main cell types between SpatialCTD healthy liver tissue sample and hepatocellular carcinoma sample. These genes are highly expressed in SpatialCTD hepatocellular carcinoma, but low expressed in SpatialCTD healthy liver tissue.

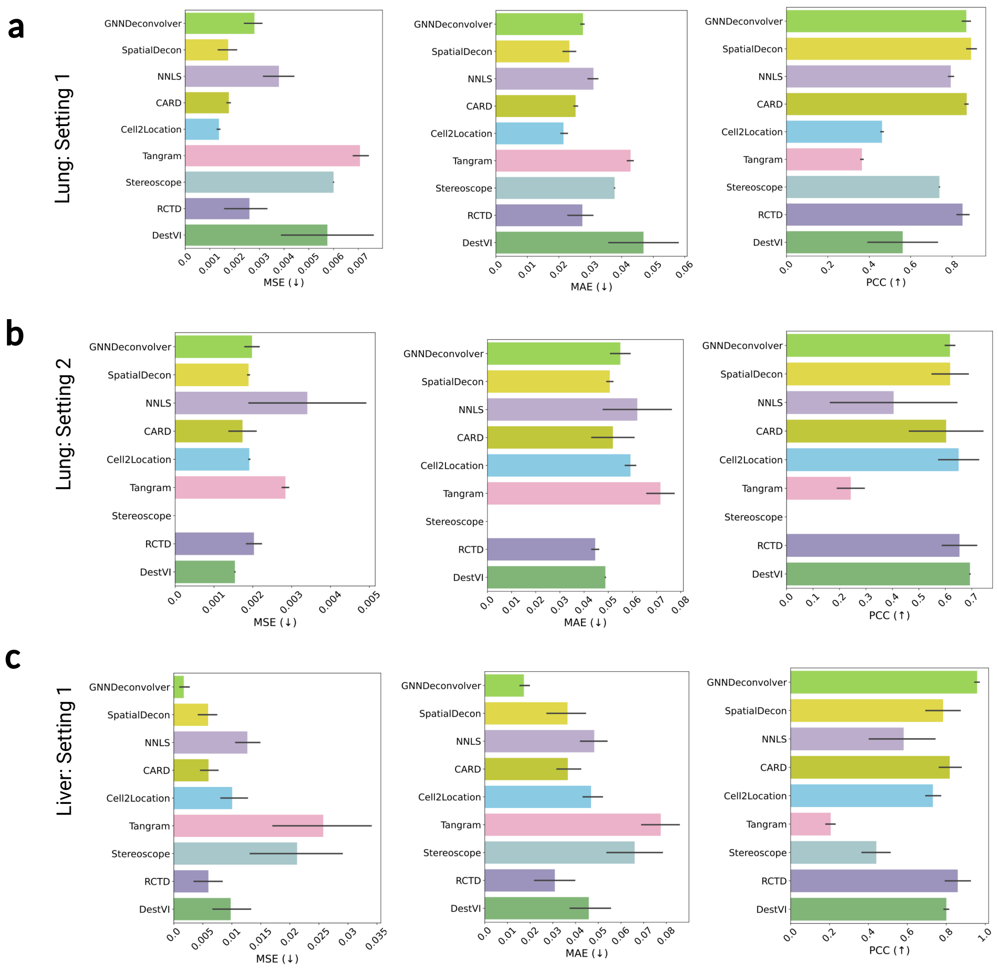


Figure 17: Performance of 9 methods in cell type deconvolution on SpatialCTD lung and liver. **a**, Comparison of the models under Setting 1 on SpatialCTD lung tissue in terms of MSE, MAE and PCC. **b**, Comparison of the models under Setting 2 on SpatialCTD lung tissue in terms of MSE, MAE and PCC. **c**, Comparison of the models under Setting 1 on SpatialCTD liver tissue in terms of MSE, MAE and PCC.



Draft
